# UBQLN2, but not an ALS-linked variant, protects client protein from temperature-induced self-association

**DOI:** 10.64898/2026.11.16.767065

**Authors:** Nirbhik Acharya, Thuy P. Dao, Nikolas Grotewold, Mallory P. Brown, Jessica K. Niblo, Maxwell B. Watkins, Jesse B. Hopkins, True Rappold, Shahar Sukenik, Stewart N. Loh, Henry L. Paulson, Lisa M. Sharkey, Carlos A. Castañeda

**Author notes:** Corresponding author: Carlos A. Castañeda. These authors contributed equally to this work (Nirbhik Acharya, Thuy P. Dao).

## Abstract

Aside from its established role in delivering ubiquitinated substrates for degradation, the proteasomal shuttle factor UBQLN2 also functions as a chaperone, likely via its central STI1 domains. Chaperone activity remains poorly understood due to the lack of known clients. Domesticated retrotransposon protein RTL8 was recently identified as a UBQLN2 interactor, and RTL8’s cellular levels are stabilized by UBQLN2. RTL8 is a small protein comprising an N-terminal helix that drives its high-order oligomerization and a C-terminal helical bundle that weakly self-associates, especially at higher temperatures. Here, we provide direct evidence that UBQLN2 acts as a chaperone for RTL8. The N-terminal region of UBQLN2, through its UBL and STI1-I domains, engages RTL8 with low-nanomolar affinity in a sandwich conformation that prevents temperature-dependent self-association of RTL8. Notably, an ALS-linked mutation in the UBQLN2 STI1-I domain reduces RTL8 affinity by an order of magnitude, and impairs the ability of UBQLN2 to protect RTL8 from self-association *in vitro* and to stabilize RTL8 levels in cells. These findings expand the functional repertoire of UBQLN2 beyond proteasomal shuttling and link its chaperone activity to a disease-relevant mutation.

**Highlights:**

- N-terminal 24 residues of retrotranspon-derived element RTL8 mediates its high-order oligomerization, while the folded C-terminal exhibits dynamic self-association
- N-terminal region of UBQLN2 (UBL and STI1-I) interacts with RTL8 in a sandwich-like mode with nanomolar affinity
- UBQLN2 chaperones RTL8 client at elevated temperature by preventing its self-association
- UBQLN2 ALS-linked mutation P189T in the STI1-I domain weakens binding to RTL8, and hinders the ability of UBQLN2 to stabilize RTL8 levels in cells

## Introduction

Ubiquilins (UBQLNs) belong to a family of ubiquitin-binding proteins best characterized for their canonical role as proteasomal shuttle factors ^1–3^. Through an N-terminal ubiquitin-like (UBL) domain that interacts with the proteasome and a C-terminal ubiquitin-associated (UBA) domain that binds ubiquitinated substrates, UBQLNs promote substrate degradation ^4–11^. Paradoxically, UBQLNs can also stabilize/protect certain clients acting as chaperones ^2,7,12–14^. The molecular determinants that govern the triage decision between degradation versus protection remain poorly understood. Malfunctioning UBQLN, via disease-linked mutations and/or dysregulation of expression levels, is linked to amyotrophic lateral sclerosis (ALS) or frontotemporal dementia (FTD) ^3,15–23^. While the roles of the UBL and UBA domains in substrate shuttling are well established, the chaperone-like activity of UBQLNs has more recently been attributed to their central STI1 region ^7,24^. This protective function is understudied due to the scarcity of characterized UBQLN clients that are neither ubiquitinated substrates nor proteasomal subunits.

A set of recently identified UBQLN2 interactors/clients includes small (∼13 kDa), nearly identical RTL8A, RTL8B and RTL8C proteins, collectively referred to as RTL8 ^25,26^. The *RTL8* family belongs to a class of genes domesticated from long terminal repeat (LTR) retrotransposons, and encodes truncated gag proteins ^27^. RTL8 retains homology to the N-terminal lobe of the retroviral capsid domain, but lacks both the C-terminal nucleic-acid-binding capsid lobe within the gag domain, and the entire pol domain which contains a protease with one active site motif for self-cleavage, as found in some related family members such as PEG10. Consistent with this truncated architecture, RTL8, when expressed alone, is not found in the fraction containing extracellular vesicles, but can be incorporated into and antagonize the formation of PEG10-derived particles ^28,29^. Relatedly, both RTL8 and UBQLN2 are required to regulate PEG10 cellular levels, the dysregulation of which is linked to ALS pathogenesis ^28–30^. This UBQLN2-RTL8-PEG10 axis is further implicated in RTL8’s role in recruitment of UBQLN2 to stress granules ^29^. On the organismal scale, knockout of *RTL8a/RTL8b* in mice leads to abnormal social behaviors and responses ^31^. Beyond these points, the intrinsic biochemical function of RTL8 remains largely undefined.

RTL8’s cellular levels positively correlate with those of UBQLN2 ^25,26^. UBQLN2’s stabilization of RTL8 is consistent with a broader chaperone-like activity ascribed to UBQLNs beyond proteasomal delivery ^7,24^. Genetic loss-of-function studies support this model directly. Proteomic profiling of *Ubqln2*-knockout mice and cells identified RTL8 among the most strongly UBQLN2-dependent proteins. Instead of promoting the degradation of clients such as PEG10 and TRIM32, UBQLN2 protects RTL8 from degradation ^25^. Notably, in the absence of UBQLN2, RTL8 protein levels are drastically reduced even as RTL8 mRNA levels increase, indicating that the loss occurs post-transcriptionally at the level of protein stability rather than expression. This dependency is specific to UBQLN2; knockout of UBQLN2, but not UBQLN1 or UBQLN4, in mice depletes RTL8 levels in brain lysates ^26^. Pulldown and colocalization experiments further showed that UBQLN2 constructs lacking UBL, PXX, or UBA domains still interact with RTL8, indicating that the interaction does not require UBQLN2’s canonical substrate-shuttling domains^26^. Despite these advances, the molecular basis of the UBQLN2:RTL8 interaction is unresolved. Recent cellular work suggested that the N-terminus of RTL8 is an important determinant of the UBQLN2 interaction, based on loss of UBQLN2 pulldown upon RTL8 N-terminal truncation ^29^. Whether this region constitutes the direct binding interface has not been established at the molecular level. Resolving these questions is also key to understanding how UBQLN2 stabilizes RTL8 and why RTL8 is cleared in its absence.

Here, we dissect the UBQLN2:RTL8C interaction at a molecular resolution using a combination of experimental and computational tools including solution nuclear magnetic resonance (NMR) spectroscopy, fluorescence anisotropy, SEC-MALS-SAXS, AlphaFold modeling, and computational simulations. We map the interaction surfaces on both UBQLN2 and RTL8C and find a novel interaction mode between them. Our data highlight that RTL8C possesses an intrinsic self-association propensity that wild-type UBQLN2, but not an ALS-associated mutant, can counteract. Together, these results provide the molecular basis, as well as a disease connection, for UBQLN2’s non-canonical, chaperone-like engagement of RTL8C.

## Results

### RTL8C is well-folded and oligomerizes in the presence of N-terminal residues 1-24

Using the AlphaFold predicted structure of RTL8C as a guide (**Fig. 1A**), we expressed and purified two recombinant RTL8C constructs from *E. coli* cells (see Methods), specifically full-length (FL) RLT8C and just residues 25-113 (RTL8C ΔN, missing the predicted N-terminal helical segment) **(Fig. S1A)**. Size exclusion chromatography - multi-angle light scattering (SEC-MALS) experiments showed that FL RTL8C exists as a polydisperse solution of high molecular weight (MDa) oligomers (**Fig. 1B**). We corroborated these observations using nuclear magnetic resonance (NMR) spectroscopy of FL RTL8C. With 1D proton (^1^H) NMR, we observed substantial peak broadening in the amide proton region, consistent with FL RTL8C tumbling slowly in solution due to a substantially larger size than expected for its molecular weight of 13 kDa (**Fig. S1B**). Together, these results indicated that FL RTL8C’s large size hinders detailed characterization of its biophysical properties. Unlike FL RTL8C, RTL8C ΔN is monomeric in solution according to SEC-MALS (**Fig. 1B**), exhibits sharp amide resonances in 1D ^1^H NMR, and displays well-dispersed peaks on ^1^H-^15^N SOFAST-HMQC NMR spectra where we monitor backbone amide resonances (**Fig. 1D**, **Fig. S1**). Therefore, we focused on RTL8C ΔN for our structural, biophysical and binding studies.

**Figure 1.**
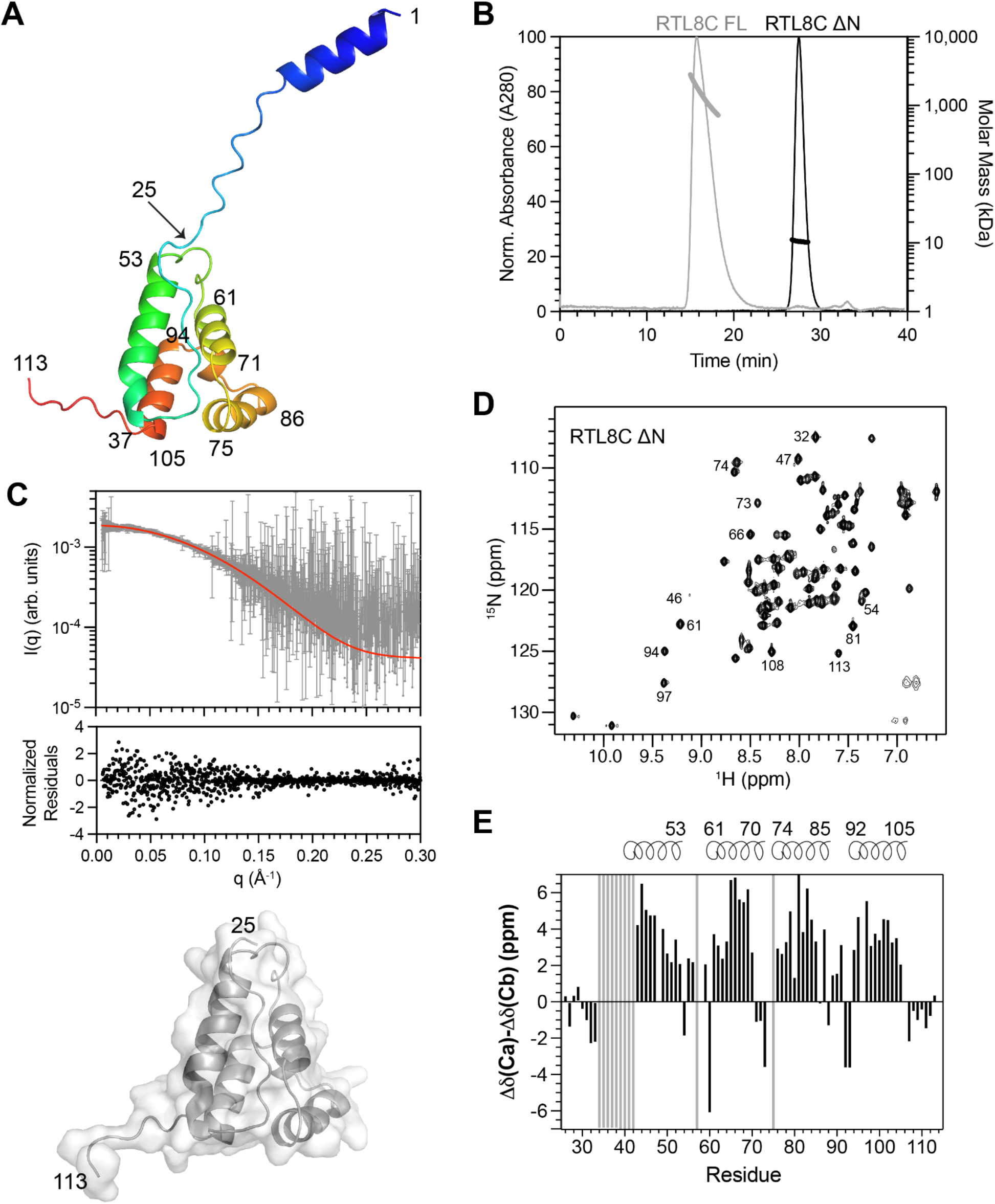
Full-length RTL8C is oligomeric while residues 25-113 (RTL8C ΔN) is a compact monomeric protein. (A) AlphaFold-predicted structure of RTL8C (AF-A6ZKI3-F1-v6) with residues rainbow-colored from blue (N-terminus) to red (C-terminus). (B) Size-exclusion chromatography (SEC) and multi-angle light scattering (MALS) of RTL8C constructs. UV absorbance at 280 nm (left axis, thin lines) is overlaid with the molar mass calculated from MALS (right axis, bold line) across the elution peak. (C) SAXS scattering curve of RTL8C ΔN with red line denoting the scattering curve from predicted single conformation of RTL8C ΔN (surface/cartoon representation shown at bottom); normalized residuals are plotted. (D) ^1^H-^15^N SOFAST-HMQC spectrum of 50 µM RTL8C ΔN at pH 7 in HEPES buffer at 25°C. (E) Residue-level secondary structure determination with experimentally determined secondary C_α_-C_β_ chemical shifts. Positive and negative ΔδC_α_-C_β_ values indicate α-helix and β-sheet content, respectively. Gray bars denote residues whose amide resonances were broadened beyond detection or not assignable.

To determine the structure of RTL8C ΔN, we compared our experimental biophysical data with that predicted from the AlphaFold structure. We obtained ^1^H, ^15^N, ^13^C backbone chemical shift assignments for 88% of RTL8C ΔN residues (see Methods). We examined secondary structure content of RTL8C using ^13^C backbone C_α_-C_β_ chemical shifts, where positive values indicate α-helical content and negative values represent β-strands (**Fig. 1E**). Our experimentally determined C_α_-C_β_ values for RTL8C ΔN correlate very well with predicted C_α_-C_β_ values from the Alphafold-derived RTL8C structure containing a four-helix bundle (**Fig. S2**; see Methods). Furthermore, the experimental small angle x-ray scattering (SAXS) data for RTL8C is in excellent agreement with the predicted SAXS scattering profile from the AlphaFold structure, as determined using a single state conformer with MultiFoXS ^32^ (**Fig. 1C**). The agreement between predicted and experimental structural data indicates that, despite the unfolding steps needed for purification (see Methods), RTL8C ΔN refolded correctly and can be used for subsequent experiments.

### RTL8C ΔN undergoes temperature-dependent self-association

Together, our SAXS and NMR data show that RTL8C ΔN is monomeric and well-folded at 25 °C. However, subtle spectral differences exist between the 50 µM and 200 µM samples resulting in both small chemical-shift perturbations (CSPs) and reduced peak intensities localized to residues 53-74 (**Fig. 2A-B**), a pattern indicative of weak self-association. To characterize this further, we monitored backbone dynamics of RTL8C ΔN on a per-residue basis via measurement of ^15^N R_1_ and R_2_ relaxation rates and steady-state ^1^H-^15^N hetNOE experiments (**Fig. 2C, 2D, S3A-B**). Together, these experiments inform on overall rotational tumbling of the protein in solution (**Table S1**), but can also detect changes in backbone dynamics on different timescales. Previously, we have used these NMR indicators to identify residues involved in self-association ^33–36^. We observed that many of the same (or nearby) residues with increased concentration-dependent CSPs also exhibit elevated R_2_ relaxation rates or lowered hetNOE values, indicative of increased backbone dynamics in the ms-µs or ns-ps timescales, respectively (**Fig. 2C, 2D, S3A**). We find that these residues (with concentration-dependent CSPs, changes in peak intensities, elevated R_2_ rates, and/or lowered hetNOE values) map to a contiguous surface on the predicted AlphaFold structure of RTL8C ΔN (**Fig. 2E**). We hypothesize that these residues comprise the surface mediating RTL8C ΔN self-association.

**Figure 2.**
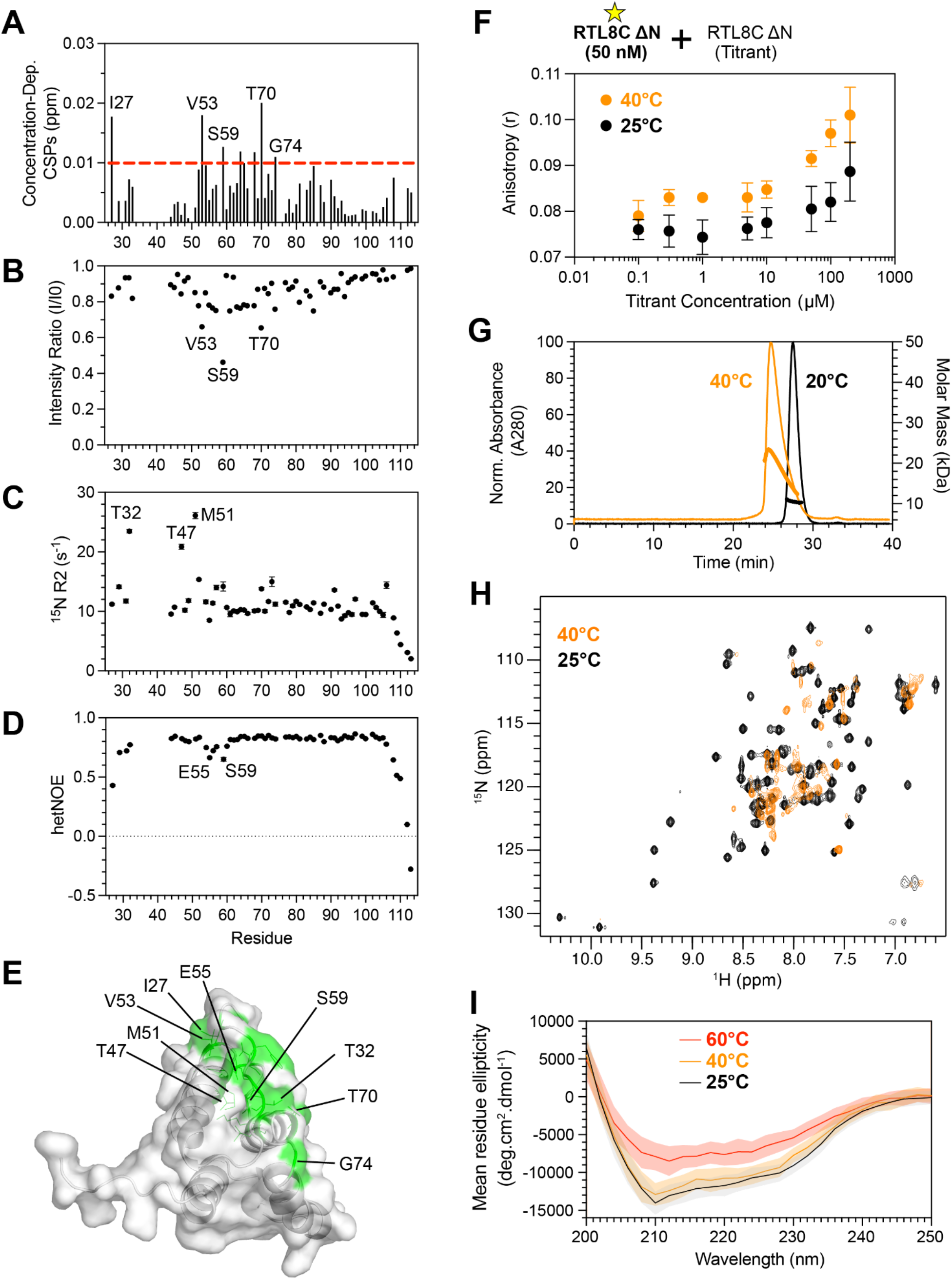
RTL8C ΔN self-associates with increasing temperature. (A) Concentration-dependent chemical shift perturbations (CSPs) and (B) peak intensity ratios of RTL8C ΔN backbone amide resonances were determined from ^1^H-^15^N SOFAST-HMQC spectra at 25 °C between 50 µM (I_0_) and 200 µM (I) RTL8C ΔN. (C) Backbone R2 relaxation rates measured using 50 µM protein. Errors were determined using 500 Monte Carlo trials using RELAXFIT. (D) hetNOE experiments were collected using 200 µM protein. Errors were determined using the standard error propagation formula. (E) Residues with concentration-dependent CSPs > 0.01 ppm, intensity ratios < 0.65, elevated R2 relaxation rates (labeled in panel C), or low hetNOE ratios (labeled in panel D) are colored green on the AlphaFold-predicted surface of RTL8C ΔN. (F) The concentration-dependent self-association of RTL8C ΔN monitored by fluorescence anisotropy experiments using 50 nM AF488-RTL8C ΔN and a titration series of unlabeled RTL8C ΔN at 25 or 40 °C. (G) SEC-MALS data collected at 20 and 40 °C. Molar mass values (bold lines) are shown across the elution peaks. (H)^1^H-^15^N SOFAST-HMQC experiments of 50 µM RTL8C ΔN at 25 and 40 °C. (I) Far-UV CD wavelength scans of RTL8C ΔN as a function of temperature. Shaded regions are instrument-reported errors at each wavelength (averaging time 30 s).

To extend this analysis over a wider concentration range, we monitored self-association by fluorescence anisotropy (FA), titrating a fixed Alexa Fluor 488-labelled RTL8C ΔN tracer (50 nM) with unlabelled RTL8C ΔN. Anisotropy, which monitors how quickly the fluorophore tumbles and reorients in solution, is influenced by the size and flexibility of the protein to which the fluorophore is attached, with higher anisotropy indicating larger size and/or decreased flexibility. At 25 °C, anisotropy increased with RTL8C ΔN concentration, particularly above ∼50 µM (**Fig. 2F**), consistent with the concentration-dependent self-association detected by NMR. The increase in anisotropy was more pronounced at 40 °C (**Fig. 2F**), indicating that self-association is enhanced at elevated temperature. This observation is supported by SEC-MALS data showing that RTL8C ΔN elutes earlier with larger species of 1-3 units at 40 °C instead of being monomeric at 20 °C (**Fig. 2G, Table S2)**. We corroborated the temperature effect orthogonally by NMR. Upon increasing temperature to 40 °C, we observed severe peak broadening of the RTL8C ΔN resonances, with many peaks no longer detectable (**Fig. 2H**), consistent with self-association, though such broadening can also arise from unfolding. To exclude the possibility of unfolding, we collected far-UV circular dichroism (CD) spectra across a range of temperatures. Spectra at 25 and 40 °C were highly similar, with minima near 210 and 222 nm characteristic of α-helical structure (**Fig. 2I, S3B**), indicating that RTL8C ΔN retains its secondary structure at 40 °C. Together, the CD, FA, and SEC-MALS data demonstrate a higher degree of self-association, rather than large-scale unfolding, that leads to the peak broadening we observe in NMR spectra at higher temperature.

### RTL8C ΔN interacts with the N-terminal 1-299 of UBQLN2

Although previous work demonstrated that UBQLN2 is an interactor of RTL8C, the binding mode between the two proteins is unknown ^25,26,29,30^. To investigate this interaction, we first monitored the FA of Alexa Fluor 488-labeled RTL8C ΔN in the presence of increasing amounts of FL UBQLN2, UBQLN2 1-299, 294-476 or 487-624 at 25 °C (**Fig. 3A**-**B**). The anisotropy increases for the samples containing FL UBQLN2 and UBQLN2 1-299, but remains unchanged for samples with UBQLN2 294-476 and 487-624. Together, these data suggest that RTL8C ΔN binds to the N-terminal half of UBQLN2. The final anisotropy values for RTL8C ΔN with FL UBQLN2 are higher than those with UBQLN2 1-299, indicating that the FL UBQLN2:RTL8C ΔN complex is larger in size, consistent with FL UBQLN2 being twice the size of 1-299 and also existing as a dimer in solution ^33^. Despite the differences in final anisotropy values, the fitted dissociation constants (K_d_) determined from the fitted single-site binding curves are nearly identical for FL UBQLN2 (6 ± 3 nM) and UBQLN2 1-299 (16 ± 10 nM) (**Fig. S4** and **Table 1**), further demonstrating that residues 1-299 of UBQLN2 comprise the main binding patch with RTL8C ΔN.

**Figure 3.**
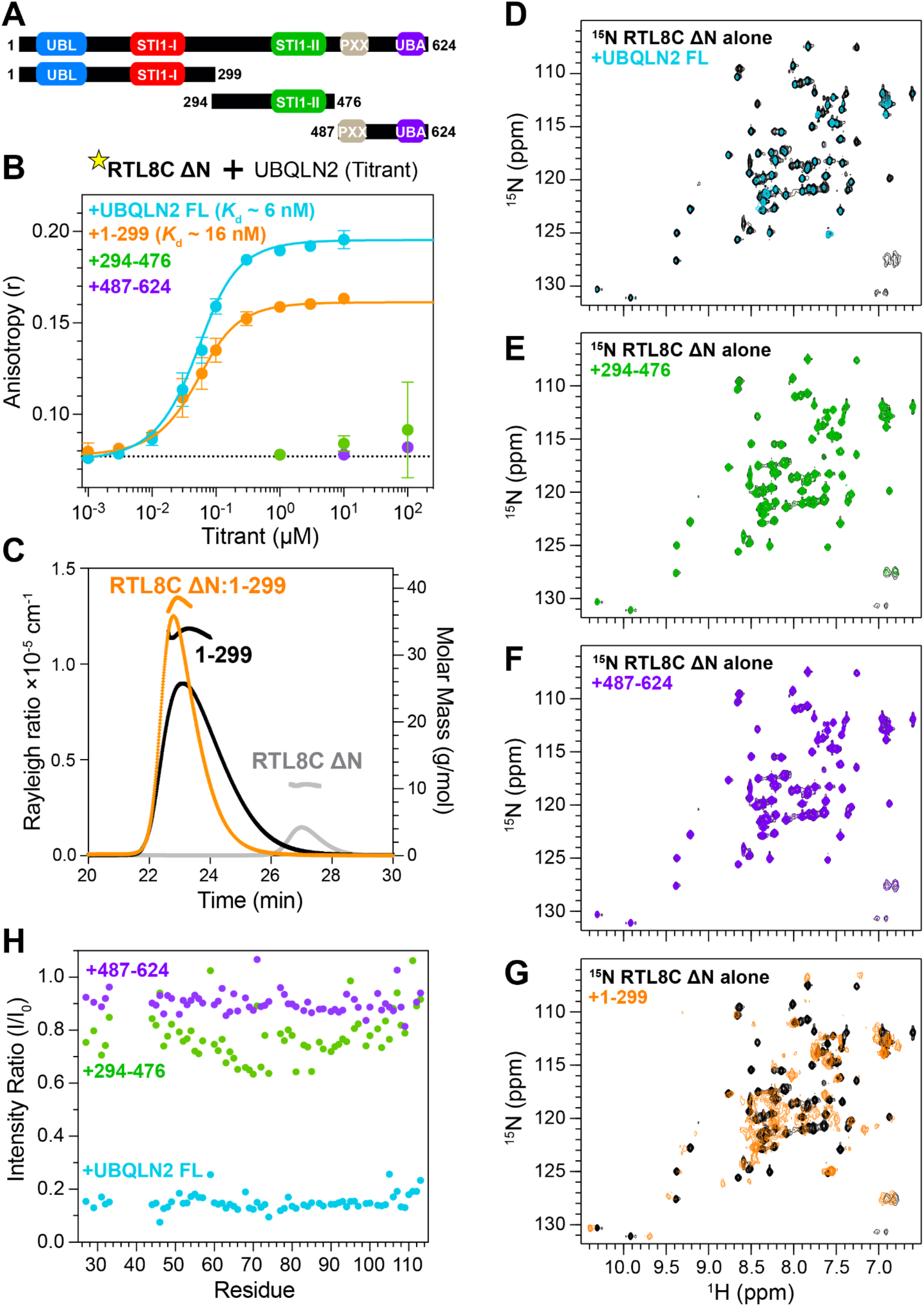
RTL8C ΔN interacts with the N-terminal half of UBQLN2. (A) Domain architecture of UBQLN2 constructs used to determine which regions bind to RTL8C ΔN. (B) Fluorescence anisotropy experiments for Alexa Fluor 488-labelled RTL8C ΔN in the presence of increasing FL UBQLN2, 1-299, 294-476 or 487-624. Data shown are the mean ± standard deviation (SD) from n≥3 independent experiments using at least two protein preps (**Table 1**). Fits shown are from using a single-site binding model. (C) SEC-MALS analysis of RTL8C ΔN, UBQLN2 1-299, and a 1:1 mixture of RTL8C ΔN:UBQLN2 1-299. Molar mass values (bold lines) are shown across the elution peaks. (D, E, F, G) ^1^H-^15^N SOFAST-HMQC NMR spectra of 50 µM RTL8C ΔN in the absence and presence of equimolar amounts of FL UBQLN2 (D), UBQLN2 294-476 (E), UBQLN2 487-624 (F) and UBQLN2 1-299 (G) at 25°C in NMR buffer (20 mM Hepes, 0.5 mM EDTA, 0.02% NaN_3_ pH 7). (H) Residue-specific intensity ratio (I/I_0_) between resonances for RTL8C ΔN in the presence (I) or absence (I_0_) of FL UBQLN2, 294-476, or 487-624.

**Table 1.** Binding affinities for RTL8C ΔN to UBQLN proteins.

|  | UBQLN2 construct (Titrant) | *K*<sub>d</sub> ± SD (25°C)<sup>a</sup> |
|---|---|---|
| RTL8C ΔN | FL | 6 ± 3 nM (n=4) |
| RTL8C ΔN | 1-299 | 16 ± 10 nM (n=4) |
| RTL8C ΔN | 1-299 P189T | 0.25 ± 0.02 µM (n=3) |
| RTL8C ΔN | 1-299 A282V | 3 ± 1 nM (n=3) |
| RTL8C ΔN | 109-299 | 0.7 ± 0.2 µM (n=3) |
| RTL8C ΔN | 109-299 P189T | 8.5 ± 1.5 µM (n=3) |
| RTL8C ΔN | 1-107 | 4.1 ± 0.3 µM (but no upper baseline) (n=3) |
| RTL8C ΔN | UBQLN1 | 0.4 ± 0.1 µM (n=3) |
| RTL8C ΔN | UBQLN4 | 68 ± 8 µM (but no upper baseline) (n=3) |
| RTL8C ΔN P75E/L77E | 1-299 | 8.5 ± 5.3 µM (but no upper baseline) (n=3) |
| RTL8C ΔN P75E/L77E | 1-107 | 2.9 ± 0.2 µM (but no upper baseline) (n=3) |

<sup>a</sup> Dissociation constants (*K*<sub>d</sub>) were determined by fluorescence anisotropy titration of Alexa Fluor (AF) 488 labelled RTL8C ΔN (WT or P75E/L77E) with the indicated UBQLN2 construct. Values are mean ± SD from n independent experiments (n as indicated).

We next performed SEC-MALS-SAXS experiments to determine the binding stoichiometry between UBQLN2 1-299 and RTL8C ΔN. MALS data report the observed molecular weight of the complex at 38 kDa, consistent with the expected molecular weight of a 1:1 complex (42.7 kDa) (**Fig. 3C**, **Table S2**). SEC data are also consistent with 1:1 binding between UBQLN2 1-299 and RTL8C ΔN, with the complex eluting as a single peak just slightly earlier than UBQLN2 1-299 despite an ∼30% increase in molecular weight. These data validate our use of the single-site binding equation for FA-derived binding assays.

To probe the interaction between RTL8C ΔN and UBQLN2 at a residue-specific level, we compared the ^1^H-^15^N SOFAST-HMQC NMR spectra of RTL8C ΔN in the absence and presence of equimolar amounts of FL UBQLN2, UBQLN2 1-299, 294-476 or 487-624 (**Fig. 3D-G**). We observed no peak movement (**Fig. 3D**) but ∼85% decrease in peak intensity (**Fig. 3H**) across RTL8C ΔN in the presence of FL UBQLN2, consistent with RTL8C ΔN interacting with FL UBQLN2 and tumbling as a much bigger complex in solution. As expected from the lack of interaction between UBQLN2 294-476 or 487-624 and RTL8C ΔN as indicated by FA experiments, minimal changes are observed on NMR spectra of RTL8C ΔN in the presence of either UBQLN2 construct (**Fig. 3E, 3F**, and **3H**). In contrast, the presence of UBQLN2 1-299, containing the UBL and STI1-I domains, resulted in substantial peak movement and/or attenuation of many RTL8C ΔN resonances (**Fig. 3G**). These data are consistent with the anisotropy data showing that RTL8C ΔN only interacts with UBQLN2 1-299. We also observed chemical shift perturbations and intensity losses to the resonances on the TROSY-HSQC NMR spectra of UBQLN2 1-299 upon RTL8C ΔN addition, confirming the interaction from the UBQLN2 side (**Fig. S5**).

### Both the N-terminal UBL and STI1-I domains of UBQLN2 interact with RTL8C ΔN in a sandwich-like conformation

As the UBL domain is a known interactor of proteasomal receptors and the STI1-I domain is believed to interact with client proteins, we hypothesized that the STI1-I domain is the main interaction driver with RTL8C ΔN. To test this, we compared the ^1^H-^15^N SOFAST-HMQC NMR spectra of RTL8C ΔN in the absence and presence of UBQLN2 constructs 1-107 (UBL) and 109-299 (STI1-I) (**Fig. 4A and 4B**). To our surprise, RTL8C ΔN resonances exhibited substantial peak movements in the presence of UBQLN2 109-299 while another subset of resonances moved in the presence of UBQLN2 1-107. To determine the RTL8C ΔN residues that interact with these two UBQLN2 constructs, we calculated CSPs (**Fig. 4A & 4B**; see Methods) and mapped them out on the AlphaFold-predicted structure for RTL8C ΔN (**Fig. 4C**). Intriguingly, UBQLN2 1-107 and 109-299 bind to RTL8C ΔN on almost opposite surfaces, with a common surface near the middle of the protein. Many of the RTL8C ΔN residues perturbed by the presence of UBQLN2 109-299 are the same ones involved in RTL8C ΔN self-association (**Fig. 4C, Fig. S6**). We confirmed the separate UBQLN2 1-107:RTL8C ΔN and UBQLN2 109-299:RTL8C ΔN interactions with FA titration experiments (**Fig. 4D, S4, Table 1**). The fitted K_d_ is lower with UBQLN2 109-299 (0.7 ± 0.2 µM) than with 1-107 (4.1 ± 0.3 µM), suggesting that UBQLN2 109-299 binds RTL8C ΔN with higher-affinity than the 1-107 region. Notably, the anisotropy signal does not plateau with the UBQLN2 1-107 titration, so the fitted K_d_ value is a lower bound estimate and additional interactions may be taking place at higher titrant concentrations.

**Figure 4.**
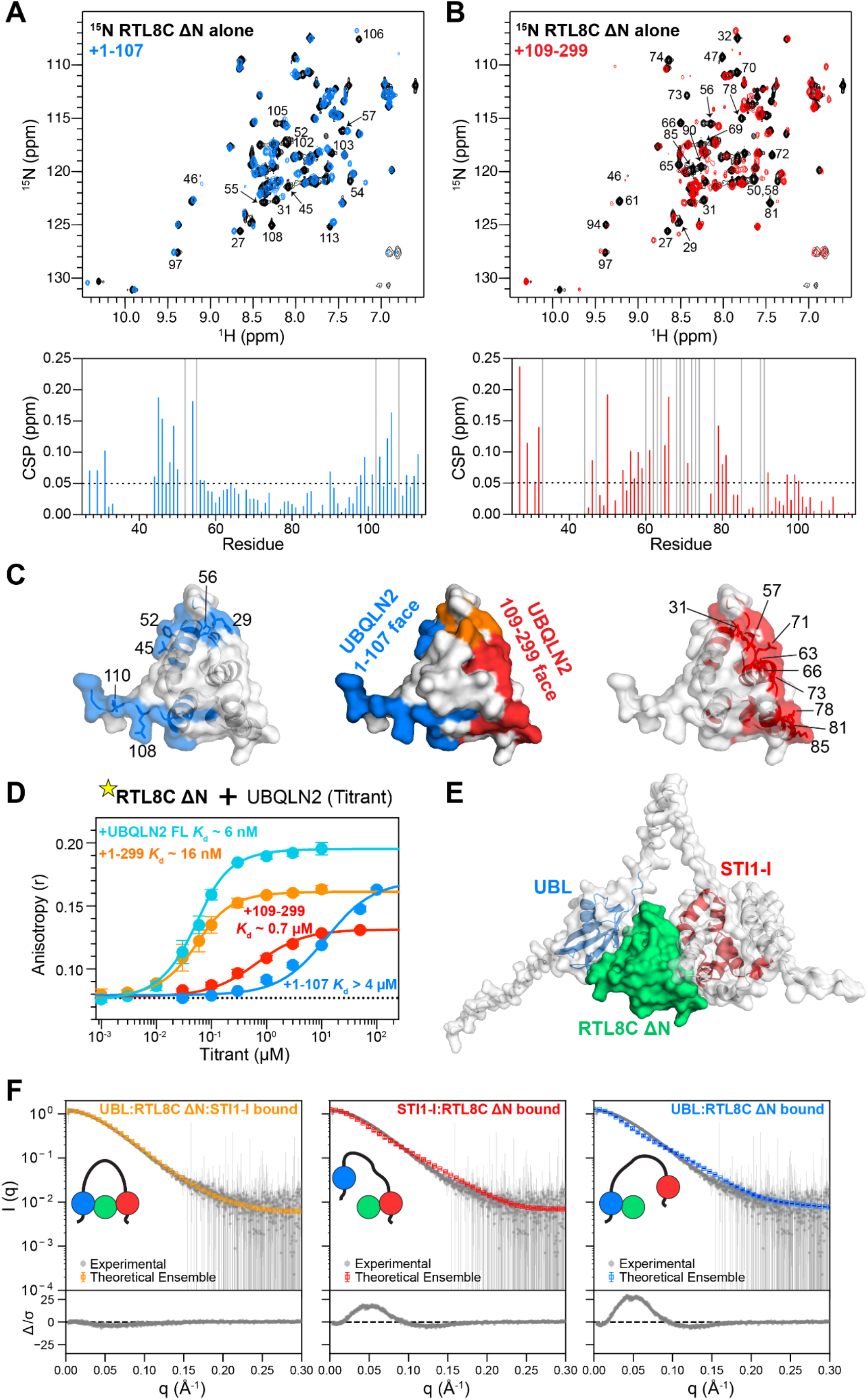
UBQLN2 UBL and STI1-I domains engage RTL8C ΔN in a 1:1 complex exhibiting a sandwich-like conformation. (A, B) ^1^H-^15^N SOFAST-HMQC NMR spectra (top) and CSP plots (bottom) comparing 50 µM RTL8C ΔN in the absence and presence of equimolar amounts of UBQLN2 1-107 (A) or UBQLN2 109-299 (B) at 25°C in pH 7 NMR buffer. NMR spectra shown with identical acquisition, processing, and visualization parameters. (C) Comparison of RTL8C ΔN binding surface with UBQLN2 constructs. RTL8C ΔN residues with CSPs > 0.05 ppm (above dotted black line in panels A and B) in the presence of UBQLN2 1-107, 109-299 or either are highlighted as blue, red or orange, respectively, that map to separate binding surfaces shown on the AlphaFold-predicted structure of RTL8C ΔN (AF-A6ZKI3-F1-v6). (D) Fluorescence anisotropy experiments for Alexa Fluor 488-labelled RTL8C ΔN in the presence of increasing FL UBQLN2, UBQLN2 1-299, 1-107 or 109-299 (see Methods). Total concentration of RTL8C ΔN was 100 nM. Data are the mean ± standard deviation (SD); n≥3 experiments using at least two protein preps. (E) Top AlphaFold-Multimer structural model of the UBQLN2 1-299:RTL8C ΔN complex, with RTL8C ΔN (green surface) sandwiched between the UBL (blue) and STI1-I (red) domains of UBQLN2 1-299 (white semitransparent surface). (F) Experimental SAXS scattering profile of UBQLN2 1-299:RTL8C ΔN complex (grey) compared against average scattering curves calculated by FoXS using simulated conformational ensembles with RTL8C ΔN bound to both UBL and STI1-I (orange; left), or only STI1-I (red; middle) or only UBL (blue; right). Normalized residuals are plotted.

The significantly lower K_d_ with UBQLN2 1-299 (16 ± 10 nM) than with either individual domain suggests that both the UBL and STI1-I regions contribute to RTL8C ΔN binding. This enhancement could reflect one of two scenarios: either two RTL8C ΔN molecules binding at separate sites, or a single RTL8C ΔN simultaneously engaging both regions. The 1:1 stoichiometry established by SEC-MALS (**Fig. 3C**) excludes the former, indicating that a single RTL8C ΔN molecule engages both the UBL and STI1-I regions simultaneously.

To visualize the engagement of RTL8C ΔN by the UBL and STI1-I domains, we obtained a structural model of the UBQLN2 1-299:RTL8C ΔN complex using AlphaFold-Multimer (**Fig. 4E**). All five predicted models positioned the RTL8C ΔN in a sandwich-like conformation between the UBL and STI1-I domains (**Fig. S7**). Notably, the interfacial residues in these models, on both RTL8C ΔN and the UBL side of UBQLN2 1-299, agreed with those independently mapped by NMR (**Fig. 4C, S5**).

To test the validity of the top UBQLN2 1-299:RTL8C ΔN AlphaFold model, we obtained the solution scattering profile of the UBQLN2 1-299:RTL8C ΔN complex from SEC-MALS-SAXS experiments. We first assessed the overall dimensions of the free and bound species, then compared the experimental scattering profile of the complex against the structural models. The experimental R_g_ of the bound complex (29.9 ± 0.1 Å) was smaller than that of free UBQLN2 1-299 (34.9 ± 0.2 Å), despite the added mass of RTL8C ΔN (R_g_ of 14.9 ± 0.2 Å) (**Table S2**). This compaction upon binding is consistent with the AlphaFold-predicted sandwich-like conformation. To compare the scattering profiles, we performed coarse-grained CALVADOS3 simulations optimized for multidomain proteins^37^ to construct a conformational ensemble of the UBQLN2 1-299:RTL8C ΔN complex. We constrained the sandwich UBL:RTL8C ΔN:STI1-I binding interfaces (as defined by the top AlphaFold model) while other segments of UBQLN2 1-299 were considered disordered (residues 1-31, 103-161, 261-299; see Methods). We calculated the average scattering curve from the conformational ensemble using FoXS ^32,38^ and simulation-derived R_g_ (29.2 ± 0.2Å, **Table S3**), and found excellent agreement with the experimental data (**Fig. 4F** left, **Movie S1**). We also tested scenarios where only the UBL:RTL8C ΔN interface or the RTL8C ΔN:STI1-I interface was preserved, but these failed to recapitulate the experimental solution scattering data and R_g_ (**Fig. 4F** center, right, **Table S3**). Together, the NMR, SAXS, and simulation data support a sandwich-like binding mode for the UBQLN2:RTL8C ΔN interaction.

### The binding between UBQLN2 and RTL8 is highly selective

To determine whether this UBQLN2:RTL8 binding mode is selective for RTL8 among related retrotransposon-derived proteins, we compared RTL8C’s sequence to its closest paralog LDOC1 (RTL7), which shares the capsid-homology fold but not RTL8’s functional role with UBQLN2 ^29^. Sequence alignment shows that the two proteins differ markedly in their N-terminal regions, but the sequence is more conserved across the C-terminal truncated gag domain (**Fig. S8A**). Even within the more conserved C-terminus, RTL8C and LDOC1 differ at several positions. Two of these positions (RTL8C Pro75 and Leu77, **Fig. S8B**) that locate to the STI1-I binding interface (**Fig. 4B, 4C**), are glutamates in LDOC1. Introducing these substitutions (P75E/L77E) into RTL8C ΔN reduced its affinity to UBQLN2 1-299 by at least 500-fold, as measured by FA (**Fig. S8C**). More broadly, the near sequence identity of RTL8 paralogs (A/B/C; **Fig. S8A**), especially across the UBQLN2 interacting interface (residue 25-113), suggests that UBQLN2’s strong affinity for RTL8C could extend to the entire RTL8 family.

We then asked whether RTL8 binds tightly to only UBQLN2 or more generally to proteins with highly conserved UBL-linker-STI1-I architecture, including paralogs UBQLN1 and UBQLN4. UBQLN2 is more closely related to UBQLN1 than UBQLN4 for both the UBL (94% vs 74% identity) and STI1-I (92% vs 80% identity) domains (**Fig. S8D**). Using FA, we found that RTL8C ΔN exhibits a 60-fold reduction and a > 10000-fold reduction in affinity to UBQLN1 and UBQLN4, respectively, compared to UBQLN2, indicating a strong preference for UBQLN2 (**Fig. S8E**). These results closely match recent cell-based studies where the reintroduction of UBQLN2 and UBQLN1, but not UBQLN4, fully and partially rescued RTL8 levels, respectively, in triple knockout (TKO) cells for UBQLN1, UBQLN2, and UBQLN4 ^26^. Together, these results indicate that the high affinity binding to UBQLN2 stabilizes RTL8C in cells.

### UBQLN2 1-299 stabilizes RTL8C ΔN at physiological temperatures via STI1-I domain

Having established that UBQLN2 1-299 clamps RTL8C ΔN in a sandwich-like conformation, we compared the RTL8C ΔN surfaces engaged in self-association and in UBQLN2 binding. NMR CSPs mapped the self-association and STI1-I-binding surfaces to overlapping regions of RTL8C ΔN (**Fig. S6**). We therefore hypothesized that UBQLN2 binding can protect RTL8C ΔN from temperature-induced self-association. Indeed, at 40 °C, whereas resonances on ^1^H-^15^N SOFAST-HMQC NMR spectra of RTL8C ΔN alone are severely broadened, those for RTL8C ΔN in the presence of UBQLN2 1-299 remain sharp and well-dispersed, indicative of a folded RTL8C ΔN in a UBQLN2:RTL8Ccomplex and not forming large oligomers (**Fig. 5A**). Temperature-dependent SEC-MALS-SAXS experiments for the RTL8C ΔN:UBQLN2 1-299 complex at 20 °C and 40 °C revealed nearly identical SEC-MALS (**Fig. 5B** right) and SAXS profiles (**Fig. S9**), unlike those observed for RTL8C ΔN alone (**Fig. 5B** left). Fluorescence anisotropy also illustrated that the size of RTL8C ΔN:UBQLN2 1-299 complex remains unchanged as a function of added RTL8C ΔN protein at both 25 °C and 40 °C (**Fig. 5C**). In contrast, addition of UBQLN2 294-476 or 487-476 did not affect RTL8C ΔN NMR spectra and RTL8C ΔN resonances remained severely broadened at 40 °C (**Fig. 5A**), further confirming that the binding and stabilization of RTL8C ΔN are highly specific to UBQLN2 1-299.

**Figure 5.**
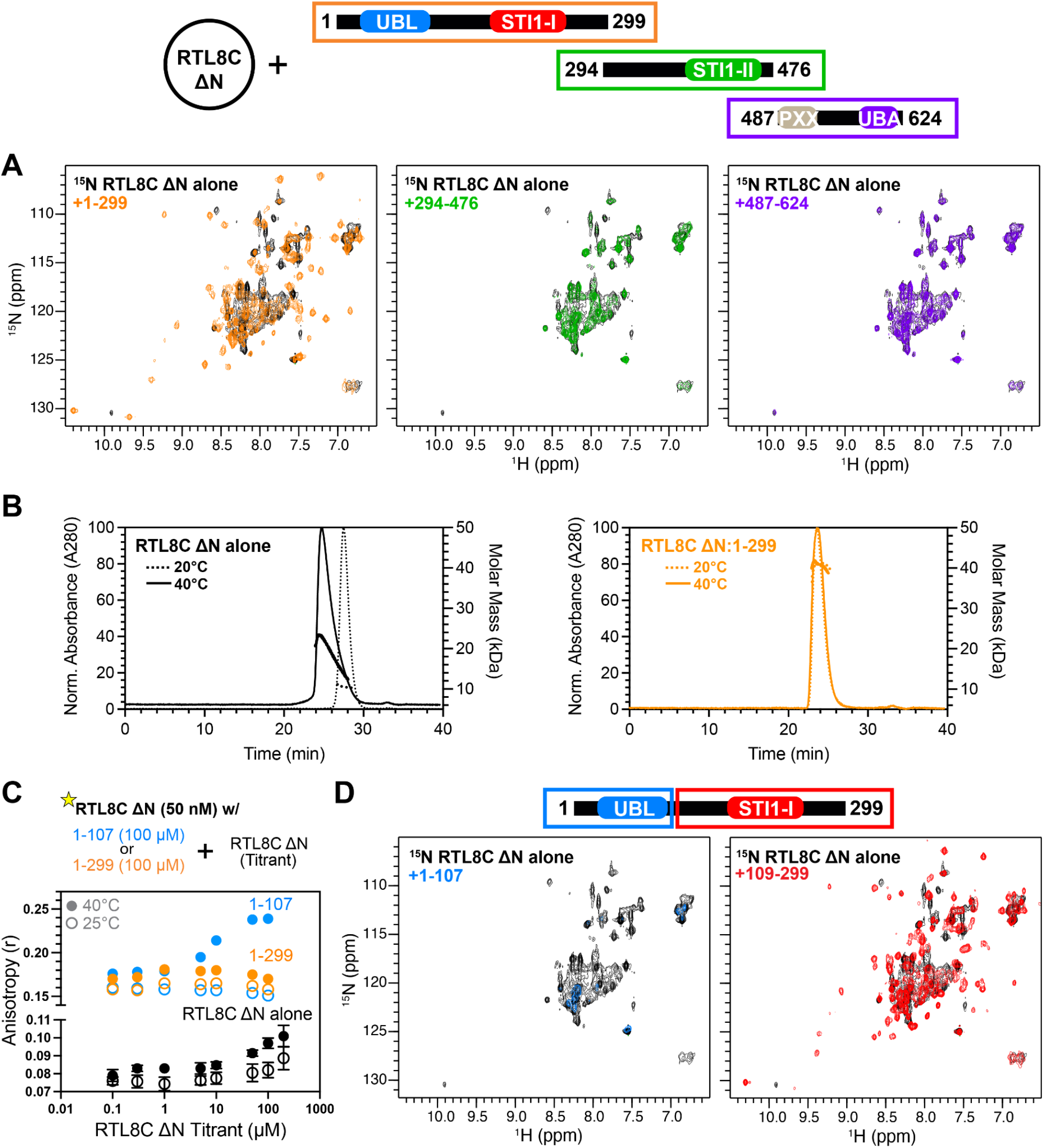
The STI1-I domain of UBQLN2 stabilizes RTL8C ΔN at physiological temperatures. (A) SOFAST-HMQC spectra of ^15^N-labeled 50 µM RTL8C ΔN with equimolar amounts of UBQLN2 constructs at 40 °C. (B) SEC-MALS data for RTL8C ΔN and 1:1 complex of RTL8C ΔN:UBQLN2 1-299 at either 20 °C or 40 °C. Molar mass values (bold lines) are shown across the elution peaks. (C) Fluorescence anisotropy experiments; schematic highlighting starting concentration of AF488 RTL8C ΔN (50 nM) and UBQLN2 construct; titration with added RTL8C ΔN. (D) SOFAST-HMQC spectra of ^15^N-labeled 50 µM RTL8C ΔN with equimolar amounts of UBQLN2 1-107 or 109-299 added at 40 °C.

If protection of RTL8C ΔN arises from occluding its self-association surface, only the UBQLN2 domain that engages that surface should be protective. Because the self-association surface overlaps the STI1-I binding interface, we predicted that UBQLN2 1-107 (containing only the UBL) would not protect RTL8C ΔN, whereas a STI1-I containing construct would. Consistent with this hypothesis, RTL8C ΔN peaks broadened extensively at 40 °C in the presence of UBQLN2 1-107 (**Fig. 5D**), indicating that RTL8C ΔN:UBQLN2 1-107 complex oligomerized even more so than RTL8C ΔN alone at 40 °C. FA experiments corroborated this, with the complex increasing in size with added RTL8C ΔN at 40 °C (**Fig. 5C**). Together, these data suggest that UBQLN2 1-107 still allows RTL8C ΔN to be prone to self-association. In contrast, RTL8C ΔN resonances in the presence of UBQLN2 109-299 are sharp and well-dispersed (**Fig. 5D**), resembling those in the presence of UBQLN2 1-299 (**Fig. 5A**). Therefore, the interaction with UBQLN2 STI1-I domain is largely, if not fully, responsible for protecting RTL8C ΔN from temperature-induced self-association.

### ALS-linked mutation in UBQLN2 hinders its ability to stabilize RTL8C

Of the three widely expressed human UBQLNs, UBQLN2 harbors the most ALS-linked mutations throughout its length ^3^. UBQLN2 109-299, which we demonstrated to be necessary to stabilize RTL8C ΔN at higher temperatures, contains four known mutations: S155N, P189T, A282V and A283T ^39,40^. To determine if these mutations affect the ability of UBQLN2 to chaperone RTL8C ΔN, we focused on P189T and A282V. In the top two AlphaFold3 models of the RTL8C ΔN:UBQLN2 1-299 complex (**Fig. 6A**), P189 is located at the interface between STI1-I and RTL8C ΔN whereas A282 is far away from any RTL8C ΔN:UBQLN2 interface and is unlikely to be involved in interactions between the two proteins. Using FA, we demonstrated that P189T causes a ∼15-fold reduction in affinity for the binding between either UBQLN2 1-299 or 109-299 and RTL8C ΔN (**Fig. 6B, Fig. S4** and **Table 1**). Therefore, P189T appears to disrupt the binding between UBQLN2 and the self-association surface of RTL8C ΔN. Unlike P189T and as expected for its predicted location, A282V had minimal effects on the binding between UBQLN2 1-299 and RTL8C ΔN (**Fig. 6B, Fig. S4** and **Table 1**).

**Figure 6.**
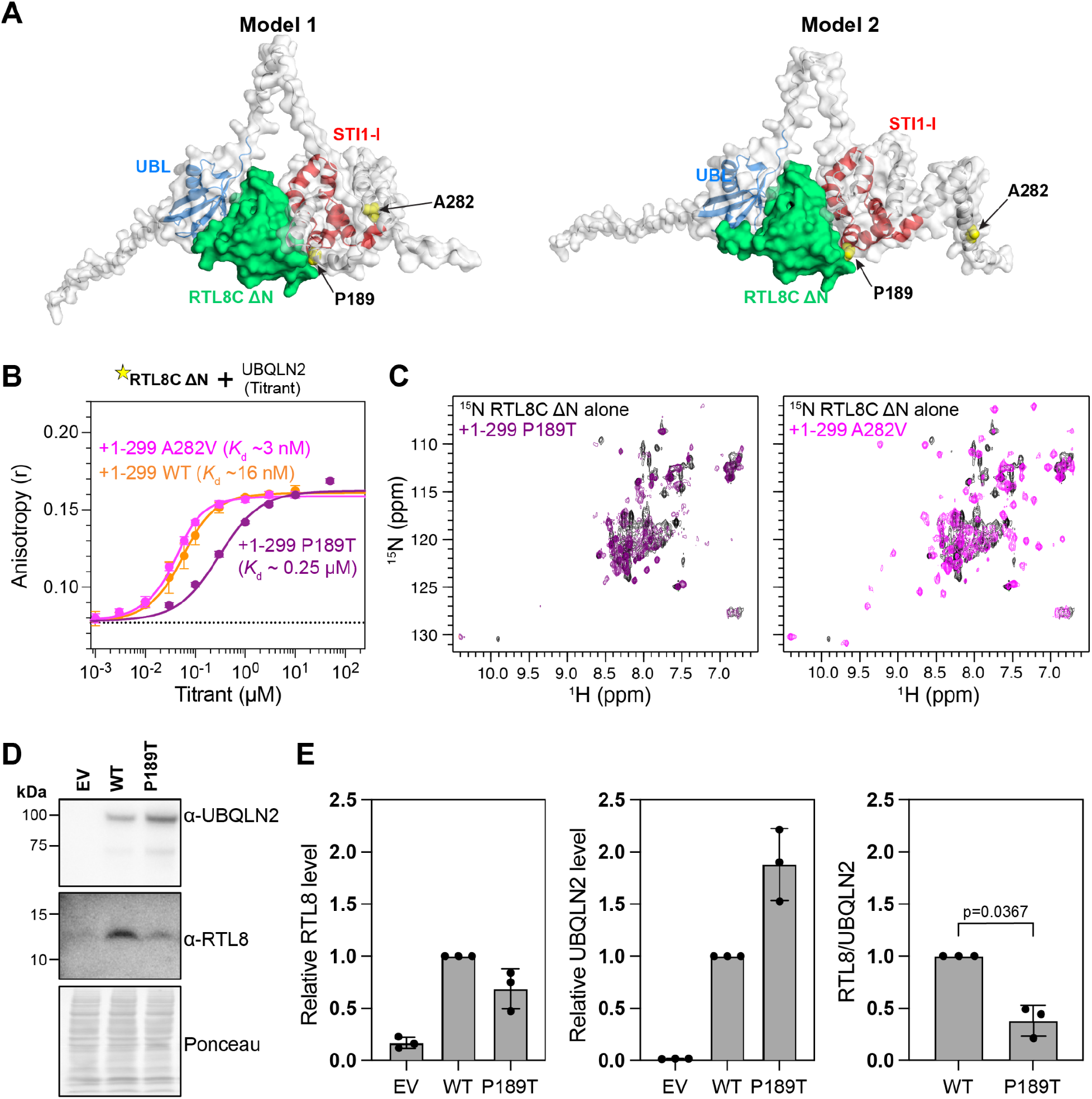
UBQLN2 ALS-linked P189T variant is unable to stabilize RTL8 due to reduced binding. (A) Location of mutation sites from the top two predicted AlphaFold3 models of RTL8C ΔN:UBQLN2 1-299 complex. Models are aligned such that RTL8C ΔN is in the same orientation. (B) Fluorescence anisotropy experiments of titrating different UBQLN2 1-299 constructs to AF488 RTL8C ΔN. Total RTL8C ΔN concentration was 100 nM. (C) SOFAST-HMQC spectra of 50 µM RTL8C ΔN at 40 °C (pH 7) in the absence and presence of equimolar amounts of P189T or A282V variant of UBQLN2 1-299. (D) Representative immunoblots for UBQLN2 and endogenous RTL8 levels in UBQLN1/2/4 triple-knockout (TKO) HEK293 cells transfected with an empty vector (EV), mEos3.2-tagged WT, or P189T UBQLN2 fusion constructs. (Bottom) Ponceau S-stained membrane showing total loaded protein. (E) Quantifications of RTL8 (left) and UBQLN2 (middle) levels shown in (D), normalized to total protein using Ponceau S staining and relative to WT UBQLN2-transfected cells within individual experiments. (Right) Ratio of RTL8 to UBQLN2 protein abundance in WT or P189T UBQLN2 transfected cells, normalized to WT UBQLN2. Data represent mean ± SD (n = 3 independent experiments) and statistical comparisons were performed using two-tailed, one-sample t-tests against 1.0, with Holm–Šídák correction (P189T, p = 0.0367).

To determine if the decrease in binding affinity to RTL8C ΔN caused by P189T affects the ability of UBQLN2 1-299 to protect RTL8 from self-association, we collected ^1^H-^15^N SOFAST-HMQC NMR spectra of RTL8C ΔN in the presence of different mutant constructs. At 25 °C, we observed similar changes in RTL8C ΔN spectra in the presence of WT UBQLN2 1-299, P189T and A282V (**Fig. S10**). At 40 °C, many resonances of RTL8C ΔN in the presence of UBQLN2 1-299 P189T are broadened or missing (**Fig. 6C** left, **Fig. S10**). In contrast, resonances for RTL8C ΔN in the presence of UBQLN2 1-299 A282V, like those in the presence of UBQLN2 1-299, remain sharp and well-dispersed (**Fig. 6C** right). Together, these data suggest that P189T, but not A282V, hinders UBQLN2’s ability to protect RTL8C ΔN from self-associating, especially at higher temperatures.

Since the P189T mutation hinders the ability of UBQLN2 to prevent the self-association of RTL8C ΔN, we hypothesized that it would also affect the ability of UBQLN2 to stabilize full-length RTL8 levels in cells. Using UBQLN TKO HEK293 cells, we transiently transfected either WT UBQLN2 or P189T UBQLN2. WT UBQLN2 restored RTL8 levels as previously observed ^26^. In contrast, P189T UBQLN2 was less efficient at rescuing RTL8 levels, especially when normalized to UBQLN2 expression levels (**Fig. 6D and 6E**). These results are also consistent with the correlation we observed between the lower affinity between UBQLN1 and RTL8 (**Fig. S8E**) and the reduced ability of UBQLN1 to rescue RTL8 levels in cells ^26^. Together, our data demonstrate that the tight and highly selective interaction between UBQLN2 and RTL8C enables the stabilization of RTL8 in cells.

## Discussion

Our work establishes the molecular basis for chaperone-like behavior of the protein quality control shuttle protein UBQLN2. UBQLN2 engages the retrotransposon protein RTL8C through a bipartite interface, sandwiching a single RTL8C molecule between its UBL and STI1-I domains with nanomolar affinity and 1:1 stoichiometry (**Fig. 3, 4**). The involvement of both the UBL and STI1-I distinguishes this interaction mode from previously proposed STI1-mediated chaperone interactions, specifically STI1-groove interactions with client helices ^36,41,42^. The surface of RTL8C that binds the STI1-I domain of UBQLN2 coincides with the same surface RTL8C uses to self-associate, and UBQLN2’s engagement suppresses temperature-induced self-association of RTL8C. UBQLN2 thus protects RTL8C by targeted occlusion of its self-association-prone surface, an interaction that is weakened by an ALS-associated mutation (P189T) in the STI1-I region of UBQLN2. Together, these results extend UBQLN2’s role beyond a proteasomal shuttle protein to a chaperone of a self-association-prone client.

Recent cellular work found that full-length RTL8C, but not an N-terminally truncated form (RTL8C ΔN), was recovered in coimmunoprecipitation (co-IP) assays with UBQLN2, implicating the N-terminus of RTL8C as the UBQLN2 binding interface ^29^. Moreover, chemical crosslinking was required to increase the amount of RTL8C in the co-IP, indicating an apparently weak interaction. These results are contradictory to our finding that RTL8C ΔN binds UBQLN2 with high affinity in the nM range (**Fig. 3**). One possible reason for the apparently weak co-IP interaction is rapid dissociation of the complex, causing substantial loss of RTL8C bound to UBQLN2 with each wash step. Indeed, we found that the complex between UBQLN2 1-299 and RTL8C ΔN is short-lived (t½ ≤ 5 s; **Fig. S11**), possibly resulting in the inability of UBQLN2 to retain this RTL8C construct throughout a co-IP assay. In contrast, an oligomeric FL RTL8C presents multiple UBQLN2-binding interfaces per particle, generating avidity that would sustain UBQLN2:RTL8C binding through wash steps and leading to its retention in co-IPs. Together, results from co-IP assays, binding affinity measurements and dissociation kinetic assays suggest that the N-terminus of RTL8C likely acts as a modulator of avidity, rather than a direct binder, to enhance the retention of tightly bound but short-lived RTL8C:UBQLN2 complexes.

A caveat of our study is the focus on the relatively weak self-association properties of RTL8C ΔN instead of the high-order oligomeric FL RTL8C which cannot yet be examined in detail (Fig. 1B, **S1B**). Our attempts to purify the N-terminal region (residues 1-24) have been unsuccessful due to its insolubility even in the presence of the solubility tag SUMO. Our data indicate that the main driver of FL RTL8C oligomerization is self-association of its N-terminus. Therefore, we propose that two types of self-association exist in FL RTL8C. The first is via the N-terminus which might form a core (**Fig. S12**) and expose the folded C-terminus (25-113) to interact with binding partners such as UBQLN2 and PEG10. The second, dissected in this study, is via the STI1-I binding surface. In the absence of UBQLN2, the second type would be enhanced, leading to even larger RTL8 oligomers that trigger a cellular signal for removal, likely via autophagy due to size. The positive correlation between the cellular levels of UBQLN2 and RTL8C supports this hypothesis ^25,26^. Therefore, our work suggests that UBQLN2 may limit RTL8C’s size due to the interaction between UBQLN2 and the C-terminal helical bundle of RLT8C (**Fig. S12**).

The ability of UBQLN2 STI1-I to bind to the self-association surface of RTL8C ΔN explains how UBQLN2 prevents the oligomerization of the C-terminal end of RTL8C (**Fig. S6**). By binding and shielding a self-association-prone surface, UBQLN2 acts as a holdase chaperone, though one that occludes a self-association surface on a folded client protein rather than sequestering an unfolded intermediate ^43^. The holdase-like behavior invites comparison with the broader family of chaperones. Reported chaperone:client dissociation constants of some of the ATP-independent chaperones span from subnanomolar to micromolar range ^43^, and the ∼6 nM affinity measured for the UBQLN2:RTL8C ΔN complex falls near the tighter end of this range, comparable to SecB and the small heat shock protein Hsp27 ^44–46^. For model clients, Hsp40, trigger factor, and SecB dissociate with k_off_ rates of roughly 10, 1, and 0.01-0.1 s⁻¹, respectively ^47,48^. The off-rate we measured is a lower bound (k_off_ ≥ 0.13 s⁻¹; **Fig. S11**), so the UBQLN2:RTL8C ΔN complex dissociates at least as fast as SecB and possibly faster. Thus, like SecB and small heat shock proteins, UBQLN2 recognizes an exposed client surface, but uniquely on a folded protein rather than an unfolded one. This provides a mechanism for the protective, non-degradative role of UBQLN function.

Our work also establishes a role for the UBL domain in UBQLN2’s chaperone activity, raising the question of how this engagement relates to UBQLN2’s known proteasomal interactions. The isolated UBQLN2 UBL binds proteasomal ubiquitin receptors with low-micromolar affinity, including Rpn10 UIM (∼1 µM) and Rpn13 Pru (∼10 µM) ^5,6^, and we estimate the isolated UBL:RTL8C ΔN interaction to be of comparable, low-micromolar affinity (**Fig. 4, Table 1**). Notably, RTL8C ΔN and these proteasomal receptors engage the same UBL surface (**Fig. S13 A-C**), and indeed we find that RTL8C ΔN and Rpn10 UIM directly compete for it (**Fig. S13D**).

Given their comparable affinities, the balance between these competing UBL:receptor and UBL:RTL8C interactions would be set not by affinity alone but also by their dissociation rates, which govern how rapidly UBQLN2 exchanges between partners, and by the STI1-I domain, which gives RTL8C a second anchor and may also engage other clients. Together, these factors could allow RTL8C binding to tune UBQLN2’s availability for proteasomal shuttling, or conversely allow proteasomal engagement to release RTL8C. Shifting the view to RTL8C itself reveals a parallel competition, as RTL8C independently antagonizes PEG10 particle assembly and recruits UBQLN2 to stress granules ^28,29^. UBQLN2 binding could therefore modulate RTL8C’s availability for these roles.

The disease relevance of UBQLN2’s protective interaction is underscored by its sensitivity to an ALS-associated mutation. The STI1-I substitution P189T weakens the UBQLN2:RTL8C interaction by an order of magnitude. Comparison of ¹H-¹⁵N spectra for wild-type and P189T UBQLN2 1-299 shows no signature of global destabilization (**Fig. S14**), so the weakened binding does not reflect unfolding of the construct. This is consistent with a recent study where the same P189T substitution disrupts client engagement without altering UBQLN2 folding ^41^. Hence, we attribute P189T’s compromised RTL8C engagement to a local perturbation of this client-binding surface. Consistent with this, P189T UBQLN2 poorly rescues RTL8 levels in cells compared to WT UBQLN2 (**Fig. 6D-E**).

Taken together, these findings show that UBQLN2 can have opposing effects on binding partners. Beyond its established role in delivering ubiquitinated substrates for degradation, UBQLN2 can also protect a client, as we show here for RTL8C, by shielding a self-association-prone surface. Because the UBQLN2 UBL domain both engages UBQLN2 at the proteasome and helps sandwich RTL8C, competition for that domain could link UBQLN2’s degradative and protective functions. The ALS mutation P189T in UBQLN2 STI1-I domain weakens UBQLN2’s chaperone/substrate protection function but does not affect the other parts of the protein involved in proteasomal degradation. UBQLN2 dysfunction may therefore extend beyond impaired degradation to a failure of client protection. More broadly, the surface-occlusion mechanism we describe may extend to other non-ubiquitinated UBQLN clients. Defining what prioritizes a client toward protection or degradation will be key to understanding how UBQLNs maintain proteostasis and how their disruption drives neurodegeneration.

## Study Limitations

Our study focused on the temperature-induced oligomerization property of RTL8C ΔN, which is mostly monomeric at room temperature. However, FL RTL8C exists as oligomers via the interactions of its N-terminus at room temperature. How FL RTL8C behaves at higher temperatures and the functional relevance of these properties still need further investigation.

## Acknowledgements

We acknowledge support from National Institutes of Health (NIH) R35 GM158070 to C.A.C., R35 NS122302 to H.L.P., and postdoctoral support to J.K.N. from the Syracuse University Vice President of Research office. Support for the Bruker 800 MHz spectrometer with TCI cryoprobe was provided by shared instrumentation NIH Grant 1S10OD012254. We appreciate support from Dr. Charlie Fry at the SUNY-ESF NMR facility. This research was performed on APS beam time award(s) (DOI(s): https://doi.org/10.46936/APS-192362/60015871 and https://doi.org/10.46936/APS-195076/60017585 from the Advanced Photon Source, a U.S. DOE Office of Science User Facility operated for the DOE Office of Science by Argonne National Laboratory under Contract No. DE-AC02-06CH11357. BioCAT was supported by grant P30 GM138395 from the National Institute of General Medical Sciences (NIGMS) of the NIH. We thank Syracuse University’s High Performance Computing cluster, OrangeGrid (NSF ACI-1341006), for computational resources. We are grateful to Drs. Alexandra Whiteley (UC Boulder), Daniel Kraut (Villanova) and Matthew Wohlever (University of Pittsburgh) and Sami Barmada (U of Michigan) for the valuable discussion on our work. The content is solely the responsibility of the authors and does not necessarily reflect the official views of NIGMS or the NIH.

## Data Availability

The datasets and computer code produced in this study are available in the following databases:

- SAXS data: SASBDB - TBD
- NMR chemical shift assignments: BMRB - TBD
- CALVADOS simulation trajectory data: https://zenodo.org/records/23427765
- Simulation analysis code: https://github.com/sukeniklab/UBQLN2-RTL8_2026

## Author Contribution

NA: Conceptualization, Data curation, Formal Analysis, Investigation, Methodology, Software, Supervision, Validation, Visualization, Writing – original draft, Writing – review & editing; TPD: Conceptualization, Data curation, Formal Analysis, Investigation, Methodology, Software, Supervision, Validation, Visualization, Writing – original draft, Writing – review & editing; NG: Conceptualization, Data curation, Formal Analysis, Investigation, Methodology, Software, Validation, Visualization, Writing – review & editing; MPB: Formal Analysis, Writing – review & editing; JKN: Data curation, Formal Analysis, Investigation, Methodology, Software, Validation, Visualization, Writing – review & editing; MBW: Data curation, Formal Analysis, Investigation, Methodology, Software, Validation, Visualization, Writing – review & editing; JBH: Data curation, Formal Analysis, Investigation, Methodology, Software, Validation, Visualization, Writing – review & editing; TR: Data curation, Formal Analysis, Investigation, Writing – review & editing; SS: Formal Analysis, Investigation, Writing – review & editing; SNL: Data curation Investigation, Methodology, Writing – review & editing; HLP: Conceptualization, Funding acquisition, Investigation, Methodology, Supervision, Writing – review & editing; LMS: Conceptualization, Investigation, Methodology, Supervision, Writing – review & editing; CAC: Conceptualization, Data curation, Formal Analysis, Funding acquisition, Investigation, Methodology, Project administration, Supervision, Writing – original draft, Writing – review & editing

## Methods

### Gene synthesis and subcloning

The genes encoding UBQLN2 constructs and hRTL8C were codon-optimized for expression in *Escherichia coli*, synthesized and cloned into pET24b (Novagen) by GenScript (NJ, USA). hRTL8C, hRTL8C 25-113 (RTL8C ΔN), UBQLN2 294-476 was subcloned behind a SUMO tag into pET24b (Novagen), UBQLN2 1-299, 109-299 and 109-260 was subcloned behind a Ub-RGRGGx5-SUMO (UbRS) tag using NEBuilder HiFi DNA assembly kit (NEB). Different mutant constructs of hRTL8C and UBQLN2 were prepared from the original plasmid using Phusion Site-Directed Mutagenesis Kit (Thermo Scientific).

The mEos3.2-UBQLN2 mammalian expression construct was derived from the mCherry-UBQLN2 plasmid described previously ^33,49^. UBQLN2 construct containing the P189T mutation was prepared from the original plasmid using Phusion Site-Directed Mutagenesis Kit (Thermo Scientific). The mEos3.2-only control construct consisted of the mEos3.2 coding sequence and linker region without any UBQLN2 sequence. All constructs were verified by whole-plasmid nanopore sequencing (Plasmidsaurus).

### Protein Expression, and Purification

UBQLN2 and the SUMO-hRTL8C constructs as well as the SUMO-, and UbRS-tagged UBQLN2 constructs were expressed in *Escherichia coli NiCo21 (DE3)* cells (NEB) in Luria-Bertani (LB) broth (Fisher BioReagents) at 37 °C and 18 °C, respectively, overnight. NMR-active (^15^N and ^13^C/^15^N labeled) protein samples were expressed in M9 minimal media as detailed in ^33^. Bacteria were pelleted and frozen before purification.

UBQLN2 was purified via a “salting out” process as described in ^33^. Briefly, cells containing overexpressed UBQLN2 were lysed in 50 mM Tris, 1 mM EDTA (GoldBio), 1 mM PMSF (GoldBio), 50 ng/µL lysozyme (GoldBio), 4 mM MgCl_2_, 25 U of Pierce universal nuclease (Thermo Scientific) (pH 8). NaCl was added to the cleared lysate to the final concentration of 0.5 M to induce phase separation. UBQLN2 droplets were pelleted and then dissolved in 20 mM Hepes (pH 7). Leftover NaCl was removed through HiTrap desalting column (GE Healthcare).

Cells containing overexpressed UbRS-UBQLN2 109-260, UbRS-UBQLN2 1-299, and UbRS-UBQLN2 109-299 and P189T or A282V mutants were lysed in 20 mM Hepes, 500 mM NaCl, 1 mM EDTA, 1 mM PMSF, 50 ng/µL lysozyme, 4 mM MgCl_2_, 25 U of Pierce universal nuclease (pH 7.5). Cell lysates were cleared and loaded onto cation exchange column (Cytiva) and washed with 20 mM HEPES, 400 mM NaCl, 1 mM EDTA, 0.02% NaN3 buffer pH 7. The column was then incubated with 20 mM HEPES, 400 mM NaCl, 1 mM EDTA, 0.02% NaN3 buffer pH 7, 1 mM TCEP and SUMO protease Ulp for 30 minutes for on-column cleavage of the UbRS tag from the UBQLN2 constructs. Cleaved UBQLN2 constructs were further purified on HiLoad 16/600™ Superdex 75 pg (Cytiva) in 20 mM sodium phosphate, pH 6.2, 0.5 mM EDTA, 0.02% sodium azide.

Cells containing overexpressed SUMO-UBQLN2 294-476 were lysed via freeze/thaw method in 20 mM sodium phosphate buffer pH 7.4, 500 mM NaCl, 25 mM imidazole, 0.5 mM EDTA, 1 mM PMSF, 1 mM MgCl_2_, and 25 U of Pierce universal nuclease. The cleared lysates were purified by Ni^2+^ affinity chromatography and eluted in 20 mM sodium phosphate buffer pH 7.4, 250 mM imidazole, 0.5 mM EDTA. The solution was concentrated to 1-2 mL. TCEP and the SUMO protease Ulp were then added to cleave the N-terminal His-SUMO tag, which was then removed via a “salting out” process as described above, but instead of NaCl, polyethylene glycol 8000 was added to 10% to induce phase separation. UBQLN2 294-476 was further purified on HiLoad™ Superdex 75 pg (Cytiva) in 20 mM sodium phosphate, pH 6.2, 0.5 mM EDTA, 0.02% sodium azide.

Cells containing overexpressed SUMO-hRTL8C, SUMO-hRTL8C 25-113 and variants were lysed in 20 mM Hepes, 1 mM EDTA, 1 mM PMSF, 50 ng/µL lysozyme, 4 mM MgCl_2_, 25 U of Pierce universal nuclease (pH 7.5). Cell lysates were cleared and the pellets were resuspended back into 20 mM Hepes, 1% Brij-35 (Thermo Scientific) (pH 7.5). Solutions were cleared and the pellets were then dissolved into 20 mM Hepes, 0.1% sarkosyl (Fisher Scientific) (pH 7.5). Solutions were cleared again and the pellets were then dissolved into 20 mM Hepes, 1% sarkosyl (pH 7.5). Sarkosyl-soluble lysates were loaded onto Ni-NTA resin (GoldBio), washed with 20 mM Hepes, 1% sarkosyl, then washed with 20 mM NaP, 25 mM imidazole, 500 mM NaCl (pH 7.4), then washed with 20 mM Hepes (pH 7.5) and eluted with 20 mM NaPhos, 250 mM imidazole (pH 7.4). DTT and EDTA were added to the purified protein solutions to a final concentration of 1 mM each. SUMO protease Ulp (Thermo Fisher) was added in a 1:100 molar ratio and solution was dialyzed into 20 mM Hepes (pH 7.5) for 3 hours to overnight. Dialyzed protein solutions were then loaded onto Ni-NTA columns; flow-through and wash (with 20 mM Hepes, 1% sarkosyl (pH 7.5)) were collected. This step was repeated twice more to separate uncleaved SUMO-hRTL8C from cleaved. Combined Ni-NTA flow-through and wash solutions were further purified via HiTrap Q HP anion exchange column (Cytiva) using 20 mM Hepes, 1% sarkosyl (pH 7.5) as the buffer. Protein was eluted via a linear gradient from 0 to 100% of 20 mM Hepes, 1 M NaCl (pH 7.5). Fractions containing mostly hRTL8C were pooled and concentrated and further purified via size exclusion chromatography (SEC) using an ENrich^TM^ SEC 70 10 x 300 column (Biorad) in 20 mM Hepes, 1% sarkosyl (pH 7.5). SEC fractions were concentrated, then exchanged into 20 mM Hepes, 6 M Urea, pH 7.5 (to remove sarksoyl), then back to 20 mM Hepes, pH 7. Protein was again loaded onto SEC 70 10 x 300 column (Biorad) in 20 mM Hepes, pH 7 to further purification and removal of residual sarkosyl and urea.

SDS-PAGE gels were performed to confirm the purity of the proteins. Purified proteins were stored at −80 °C.

### SEC-MALS-SAXS data collection and analysis

SAXS was performed at BioCAT (beamline 18ID at the Advanced Photon Source, Chicago) with in-line size exclusion chromatography to separate sample from aggregates and other contaminants thus ensuring optimal sample quality and multiangle light scattering (SEC-MALS), dynamic light scattering (DLS) and refractive index measurement (RI) for additional biophysical characterization (SEC-MALS-SAXS). The samples were loaded onto a Superdex 200 Increase 10/300 GL column (Cytiva) run by 1260 Infinity II HPLC (Agilent Technologies) at 0.6 ml/min. The flow passed through (in order) the Agilent UV detector, a MALS detector and a DLS detector (DAWN Helios II, Wyatt Technologies), and an RI detector (Optilab T-rEX, Wyatt). The flow then went through the SAXS flow cell. The flow cell consists of a 1.0 mm ID quartz capillary with ∼20 µm walls. A coflowing buffer sheath is used to separate the sample from the capillary walls, helping prevent radiation damage ^50^. SAXS scattering intensity was recorded using a Pilatus3 × 1 M (Dectris) detector which was placed 3.6 m from the sample giving access to a q-range of 0.003 Å^-1^ to 0.33 Å^-1^. A 0.5 s exposure was acquired every 2 s during elution, and data were reduced using BioXTAS RAW 2.4 ^51^. Buffer blanks were created by averaging regions flanking the elution peak and subtracted from exposure selected from the elution peak to create the I(q) vs. q curves used for subsequent analysis. Molecular weights and hydrodynamic radii were calculated from the MALS and DLS data respectively using ASTRA 8 software (Waters). Additionally, R_g_ and I(0) values were obtained using the entire q-range of the data by calculating the distance distribution functions, P(r) vs. r, using GNOM ^52^.

In the UBQLN2 1-299:RTL8C complex SAXS sample, based on the experimental K_d_ (**Table 1**) and concentrations used for the SEC-MALS-SAXS experiment (**Table S4**), the calculated unbound fractions of RTL8 and UBQLN2 1-299 are <3%. All SEC-MALS-SAXS parameters for data collection and analysis can be found in **Table S4**.

### Comparison between RTL8C ΔN experimental and predicted scattering curves

Comparison between the experimental SAXS scattering curve and the predicted scattering curve for RTL8C ΔN (**Fig. 1**) was done using MultiFoXS ^32^ via the https://modbase.compbio.ucsf.edu/multifoxs/ site. The AlphaFold2-predicted structure of RTL8C (AF-A6ZKI3-F1) was used while defining residues 107-113 to be flexible. A total of 10,000 conformers were generated and best-fit results were determined using an ensemble containing a single state.

### NMR experiments

All NMR experiments were performed at 25-45**°**C, as specified, on a Bruker Avance III 800 MHz spectrometer equipped with TCI cryoprobe. Proteins were prepared in the NMR buffer: 20mM sodium phosphate buffer (pH 6.2), 0.5mM EDTA, 0.02% NaN_3_, and 5% D_2_O or 20 mM Hepes buffer (pH 7), 0.5mM EDTA, 0.02% NaN_3_, and 5% D_2_O. All NMR data were processed on NMR Box ^53^ using NMRPipe ^54^ and analyzed using CCPNMR 2.5.2 ^55^.

### NMR 2D SOFAST-HMQC and TROSY spectra

^1^H-^15^N SOFAST-HMQC experiments were acquired using spectral widths of 12 and 28 ppm in direct ^1^H and indirect ^15^N dimensions, respectively, and corresponding acquisition times of 80 ms and 28 ms. All SOFAST-HMQC spectra were collected with 48 scans. ^1^H-^15^N TROSY-HSQC experiments were acquired using spectral widths of 15 and 27 ppm in the direct ^1^H and indirect ^15^N dimensions, and corresponding acquisition times of 200 ms and 46 ms. Centers of frequency axes were ∼4.7 and 117 ppm for ^1^H and ^15^N dimensions, respectively. ^1^H-^15^N TROSY spectra were processed and apodized using a Lorentz-to-Gauss window function with 15 Hz line sharpening and 20 Hz line broadening in the ^1^H dimension, while ^15^N dimension was processed using a cosine squared bell function. All TROSY spectra were collected with 16 scans. Contour level and peak intensities were normalized to factor in concentration difference for fair and quantitative comparison across different concentrations wherever applicable. Chemical shift perturbations (CSPs) were quantified as follows: Δδ = [(Δδ_H_)^2^ + (Δδ_N_/5)^2^]^1/2^ where Δδ_H_ and Δδ_N_ are the differences in ^1^H and ^15^N chemical shifts, respectively.

### NMR chemical shift assignments of RTL8C ΔN

We determined backbone ^1^H, ^13^C, ^15^N assignments using traditional ^1^H-detect triple-resonance experiments (HN(CA)N, HN(COCA)N, HNCACB, CBCA(CO)NH) on ^13^C/^15^N samples containing 250 µM RTL8C 25-113 in pH 7 20 mM HEPES buffer at 25°C. All experiments used optimized parameter sets incorporating non-uniform sampling.

Acquisition times were 20 ms, 6 ms, and 75-80 ms, in the indirect ^15^N dimensions, indirect ^13^Cα/Cβ dimensions, and direct ^1^H dimensions, respectively. Spectral widths were generally 28 ppm in indirect ^15^N and 70 ppm in indirect ^13^Cα/Cβ dimensions. Experiments were acquired with 12%–25% sampling using the Poisson Gap sampling method ^56^. Spectra were processed using NMRPipe and employed standard apodization parameters and linear prediction in the indirect dimensions, and analyzed using CCPNMR on NMRBox. Using these experiments, we successfully assigned backbone resonances (H, N, C_α_, C_β_) for ∼88% of all residues.

### Secondary Structure Determination from NMR Data

For C_α_ and C_β_ secondary shift calculations, random coil chemical shifts for RTL8C were determined at https://spin.niddk.nih.gov/bax/nmrserver/Poulsen_rc_CS/ using default parameters at 25°C sample temperature and pH 7 ^57^. The predicted Δδ(C_α_-C_β_) values of RTL8C, presented in **Fig. S2**, were obtained using the EFG-CS web server (https://biosig.lab.uq.edu.au/efg_cs/) ^58^. This web server employs machine learning-based models to predict chemical shifts from protein sequences and structures, including experimental structures and AlphaFold2 predictions. For this study, we used the AlphaFold2-predicted structure of RTL8C (AF-A6ZKI3-F1) as input to generate the predicted C_α_ and C_β_ chemical shifts.

### 15N Relaxation experiments

Longitudinal (R_1_), transverse (R_2_) backbone ^15^N relaxation rates, and {^1^H}-^15^N steady-state heteronuclear Overhauser enhancement (hetNOE) were measured for RTL8C ΔN using established interleaved relaxation experiments and protocols ^59,60^. Protein concentrations used were either 50 µM or 200 µM (hetNOE). Relaxation inversion recovery periods for R_1_ experiments were 4 ms (x 2), 600 ms (x 2), and 900 ms (x 2) for RTL8C, using an interscan delay of 2.5 s. Total spin-echo durations for R2 experiments were 8 ms (x 2), 48 ms (x 2), 64 ms (x 2), 96 ms, and 120 ms (x 2) using an interscan delay of 2.5 s. Heteronuclear NOE experiments were acquired with an interscan delay of 4.5 s. All relaxation experiments were acquired using spectral widths of 12 and 28 ppm in the ^1^H and ^15^N dimensions, respectively, with corresponding acquisition times of 75 and 25 ms. Spectra were processed using squared cosine bell apodization in both ^1^H and ^15^N dimensions. Relaxation rates were derived by fitting peak heights to a mono-exponential decay using RELAXFIT ^61^. Errors in R_1_ and R_2_ were determined using 500 Monte Carlo trials using RELAXFIT on MATLAB. Errors in hetNOE measurements were determined using the standard error (SE) propagation formula.

### Rotational Diffusion Tensor Determination for RTL8C ΔN

The overall rotational diffusion tensor was determined using the program ROTDIF ^62,63^ and only residues in rigid secondary structure elements. We used relaxation data for residues 44, 45, 48, 61-66, 68, 69, 71, 77-80, 83, 84, 94-96, 99, 100, 104, and 105. The ratio of relaxation rates, ρ, was determined for each residue as ρ = (2(R_2_’/R_1_’) – 1)^-1^, where R_1_’ and R_2_’ are modified R_1_ and R_2_ values with the high-frequency contributions subtracted ^64^.

### AlphaFold2 multimer modeling for predicting interactions between the RTL8C and UBQLN2 1-299

We used AlphaFold3 ^65^ to model how UBQLN2 1-299 interacts with RTL8C ΔN. Each multimer run resulted in five predicted models ranked 1 to 5. All five models from each multimer run are shown in **Fig. S7**. In the main text, the top or the top two models of UBQLN2 1-299 interacting with RTL8C ΔN are used in **Fig. 4 and 6**, respectively.

### Circular dichroism experiments

Far-UV circular dichroism (CD) data were acquired on an Aviv Model 420 CD spectrometer. Wavelength spectra of 10 µM RTL8C ΔN were recorded using a 1 mm pathlength cuvette at 25, 40, and 60 °C scanning from 250 to 200 nm in 2 nm steps with a 30 s averaging time per point. Separately, secondary structure content of 5 µM RTL8C ΔN was monitored as a function of temperature in a 10 mm pathlength cuvette, recording CD signal at 224 nm from 25 to 65 °C in 2 °C steps, with 12 s equilibration per step, and a 15 s averaging time per measurement. A 1 nm bandwidth was used for all measurements. Raw ellipticity (mdeg) was converted to mean residue ellipticity using protein concentration, number of residues, and cuvette pathlength.

### Fluorescence anisotropy experiments

Fluorescence anisotropy (FA) was used to monitor binding between AF488-labeled RTL8C ΔN (WT or P75E/L77E) and titrations of unlabeled UBQLN2 variants. Labeled protein was prepared by reacting purified RTL8C ΔN with AlexaFluor 488 NHS ester (Thermo Fisher) at a 10:1 dye:protein molar ratio in 20 mM HEPES pH 7.0 for 1 h at room temperature, followed by SEC purification (Bio-Rad ENrich SEC 70) in the same buffer to remove free dye. The degree of labeling was 1.4 (WT) or 0.74 (P75E/L77E) as determined by absorbance at 280 and 495 nm, using the manufacturer’s correction factor for AlexaFluor 488 at A280.

All samples were prepared in 20 mM NaP pH 6.8, 0.5 mM EDTA, 0.01% Tween-20 in black-bottom 96-well plates (Greiner Bio-One) at a final volume of 100 µL per well. Labeled RTL8C ΔN was held constant, and UBQLN2 variants were titrated, with one well for each variant containing labeled RTL8C ΔN alone serving as common unbound reference. Samples were equilibrated at 25°C for 10 min prior to data acquisition. FA was measured on a SpectraMax i3x plate reader equipped with the FP-Fluor module (excitation 485 nm, emission 535 nm) pre-equilibrated to 25°C. For each well, parallel and perpendicular intensity components (I_para_ and I_perp_) were recorded. FA values (r) were calculated from the standard expression:

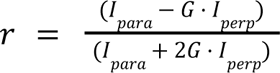

where G is the instrument’s polarization-correction factor. The G-factor was determined (G = 1.99) using free AF488 dye and applied to all FA measurements. Anisotropy titration data were fit to a quadratic binding equation that accounts for ligand depletion. The fraction of the bound complex, f_b_, was obtained from the exact solution to the 1:1 binding equilibrium:

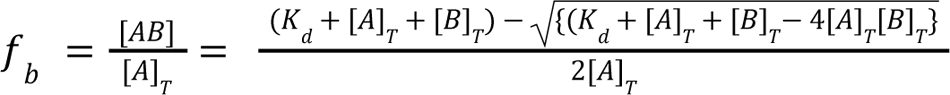

where [A]_T_ = 100 nM is the constant total concentration of RTL8C ΔN (50 nM AF488-labeled and 50 nM unlabeled) and [B]_T_ is the titrated UBQLN2 construct concentration. As the AF488 RTL8C ΔN quantum yield differs between the free and bound states for some of the titrants, anisotropy was modeled as the intensity-weighted average of the two states:

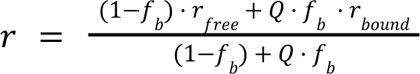

where Q = I_bound_/I_free_ is the ratio of total fluorescence intensities (I_para_ + 2G · I_perp_) in the bound versus free state averaged across replicates. Nonlinear least-squares fitting was performed in Python using the lmfit package, with K_d_, r_free_, and r_bound_ treated as free parameters bounded to physically reasonable ranges (K_d_ > 0; r values constrained to 0-0.4). Each replicate titration was fit independently, and the reported K_d_ for each construct is the mean ± SD of the K_d_ values obtained across independent replicates (UBQLN2 FL n=4; 1-299 n=4; all other constructs n=3; **Table 1**).

The concentration-dependent self-association of RTL8C ΔN and its temperature dependence, alone and in the presence of UBQLN2 constructs, were assessed by FA. Sample sets contained 50 nM AF488-labeled RTL8C ΔN and a titration series of unlabeled RTL8C ΔN, prepared either alone or with a UBQLN2 construct (1-107 or 1-299) held at a fixed concentration of 100 µM. Samples were made in 20 mM NaP pH 6.8, 0.5 mM EDTA, 0.01% Tween-20 in PCR strips and 100 µL from each was transferred to a black-bottom 96-well plate. FA was measured on a SpectraMax i3x plate reader using the FP-Fluor module (excitation 485 nm, emission 535 nm) pre-equilibrated to 25 °C. Samples were then returned to PCR strips, incubated at 40 °C for 2 h in a thermocycler, transferred (100 µL) to fresh wells of a 96-well plate, and re-measured on the plate reader pre-equilibrated to 40 °C. (RTL8C ΔN alone n = 4; with UBQLN2 constructs n = 1)

To determine the dissociation rate constant (k_off_) of the RTL8C ΔN:UBQLN2 1-299 complex, we performed a competition chase assay (**Fig. S11**). AF488 labeled RTL8C ΔN (50 nM) was pre-incubated with UBQLN2 1-299 (300 nM; >15-fold molar excess of K_d_) for 30 min at room temperature. Anisotropy of the bound complex was recorded, followed by the addition of unlabeled RTL8C ΔN as chase (15 µM; 50-fold excess over UBQLN2) to outcompete rebinding of the dissociated labeled RTL8C ΔN. Anisotropy kinetics was recorded immediately after chase addition (dead time 20-28 s). As the instrument’s actual per-point acquisition interval exceeded the nominal programmed interval (due to sequential parallel/perpendicular polarization reads), recorded time points were rescaled by the ratio of total measured run duration (6 min 32 s) to nominal run duration (5 min). Corrected anisotropy decay, anchored to the pre-chase anisotropy at t = 0, were fit to a single-exponential decay model by nonlinear least-squares regression. Free AF488-RTL8C ΔN (unbound; 50 nM) and the bound complex (with 300 nM UBQLN2 1-299) served as separate controls defining the dynamic range.

### CALVADOS coarse-grained simulations

To investigate the interactions between RTL8C and UBQLN2, we performed coarse-grained simulations using CALVADOS3_COM_. All interactions were assigned as described in the original CALVADOS3_COM_ framework ^37^, and elastic constraints were applied to folded domains as defined in **Table S3** to maintain their folded structure. The definitions of the folded domains relied on previous definitions of the UBL (residues 32-102), and the predicted STI1-I domain from AlphaFold (residues 162-260). To ensure certain folded domains remained in contact with one another, additional constraints were applied to folded domains as pairs (**Table S3**). Using the AlphaFold predicted structures ^66^, five all-atom initial conformations were generated using Modeller ^67^, which were then mapped to the CALVADOS3_COM_ representation. Simulations were run at a temperature of 298.15K, pH of 7.2, and ionic strength of 0.1M. Simulations were performed for 2 µs using Langevin dynamics, where the first 100ns were discarded as equilibration. The drag coefficient was set to 0.01 ps^−1^, and the timestep was set to 10 fs.

To enable comparison between simulations and the experimental SAXS profiles, theoretical SAXS curves were calculated using FoXS ^38^. For each simulation replicate, all 190,000 frames from its trajectory were independently passed to FoXS in residue mode to get a per-frame scattering profile, which were then averaged into a replicate profile. The resulting profiles were then averaged to give the theoretical ensemble, with the standard deviation taken as its uncertainty. Normalized residuals were calculated as (*I_exp_ - I_theoretical_)/σ_exp_*, where *I_theoretical_* is the averaged ensemble for each q-point.

### Cell culture and transient transfection

HEK293 UBQLN1/2/4 triple-knockout (TKO) cells were kindly provided by Dr. Ramanujan Hegde (Medical Research Council (MRC) Laboratory of Molecular Biology)^7^ and maintained in DMEM (Gibco™ DMEM + L-glutamine, +4.5 g/L D-glucose, + 110 mg/L sodium pyruvate) supplemented with 10% fetal bovine serum (Fisher). Cells were maintained and passaged at 5% CO_2_.

Plasmids (mEOS3.2-WT_UBQLN2, mEOS3.2-P189T_UBQLN2) were transfected into cells using Lipofectamine™ 3000 Reagent (Invitrogen) according to the manufacturer’s protocol. The P3000 reagent, DNA, OptiMEM Reduced Serum Medium (Gibco) mix was prepared separately from the Lipofectamine reagent, OptiMEM mix, prior to combining and incubating for 15 min at room temperature. The Lipofectamine/DNA/P3000 mix was then added to cells and allowed to transfect/express for 48 h.

### Immunoblotting

Protein samples prepared in 1x Laemmli buffer containing 100 mM DTT were separated by SDS-PAGE using polyacrylamide NuPAGE 4–12% Bis-Tris gradient gels (Invitrogen). Proteins were then transferred onto 0.2 μm nitrocellulose membranes at 110 V for 1 h. Membranes were rinsed with dH_2_O, stained with Ponceau S for 10 min, and imaged on a G Box Mini imager (Syngene). Membranes were then rinsed with dH_2_O and blocked using 5% non-fat dry milk (Research Products International) containing 0.05% BSA (Fisher) and 0.04% thimerosal in 1× TBST for 1 h. Membranes were incubated with primary antibodies overnight at 4°C in blocking buffer. Membranes were rinsed for 5 min with 1x TBST three times, incubated at RT with secondary antibodies in blocking buffer for 1 h, and rinsed in again three times in 1x TBST. Membranes were developed with EcoBright Nano/Femto HRP 50 (Innovative Solutions) using the G Box Mini imager set to ECL auto exposure. Blots were quantified using GeneSys (Syngene) and raw values were normalized to Ponceau S values.

| Antibody | Source | Dilution IB – immunoblotting |
|---|---|---|
| Mouse monoclonal α-UBQLN2 6H9 | Novus Biologicals #NBP2-25164 | 1:2000 (IB) |
| Rabbit polyclonal α-FAM127B/RTL8A | Proteintech #20282-1-AP | 1:500 (IB) |
| Goat α-mouse IgG(H + L) HRP-conjugated | Jackson ImmunoResearch #115-035-146 | 1:10,000 (IB) |
| Goat α-rabbit IgG(H + L) HRP-conjugated | Jackson ImmunoResearch #111-035-144 | 1:10,000 (IB) |

## Statistics

IB statistical comparisons were performed using two-tailed, one-sample Student’s t-tests against a hypothetical mean of 1.0 (normalized wild-type UBQLN2 control), adjusting P-values for multiple comparisons using the Holm–Šídák method. Statistical significance was defined as p < 0.05.

