## Supplementary Info for "UBQLN2, but not an ALS-linked variant, protects client protein from temperature-induced self-association"

**Supporting Information**

A

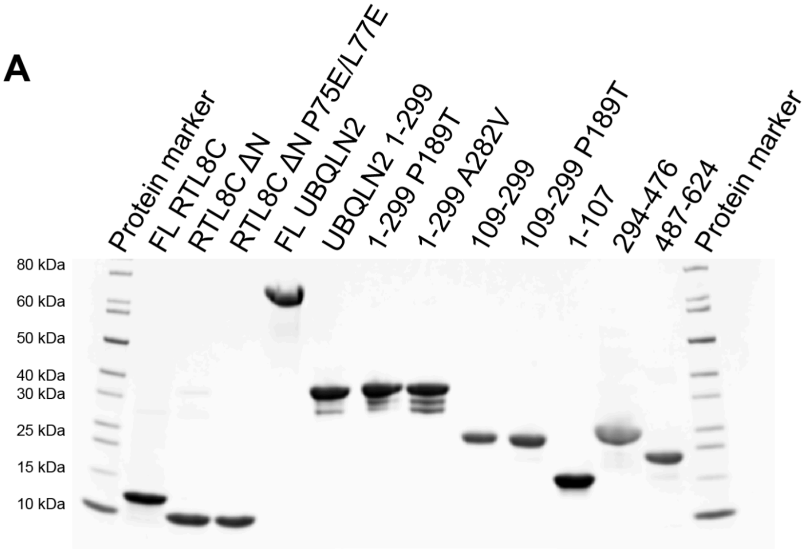


B

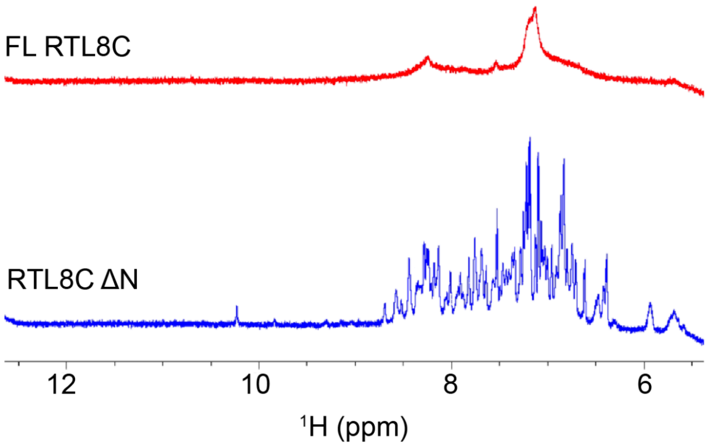


**Figure S1. Full length RTL8C shows severe peak broadening relative to RTL8C ΔN.** (A) SDS-PAGE gel of constructs used in this study. For the UBQLN2 1-299 constructs, the slightly lower molecular weight species are C-terminally truncated constructs with cleavage sites post the STI1-I domain and therefore, binding to RTL8 ΔN should not be affected. (B) 1D $^1$H NMR spectra of FL RTL8C (red) and RTL8C ΔN (blue) are compared. The amide proton region (6.5-9.5 ppm) of FL RTL8C shows substantial peak broadening, likely reflecting self-association/higher-order oligomerization.

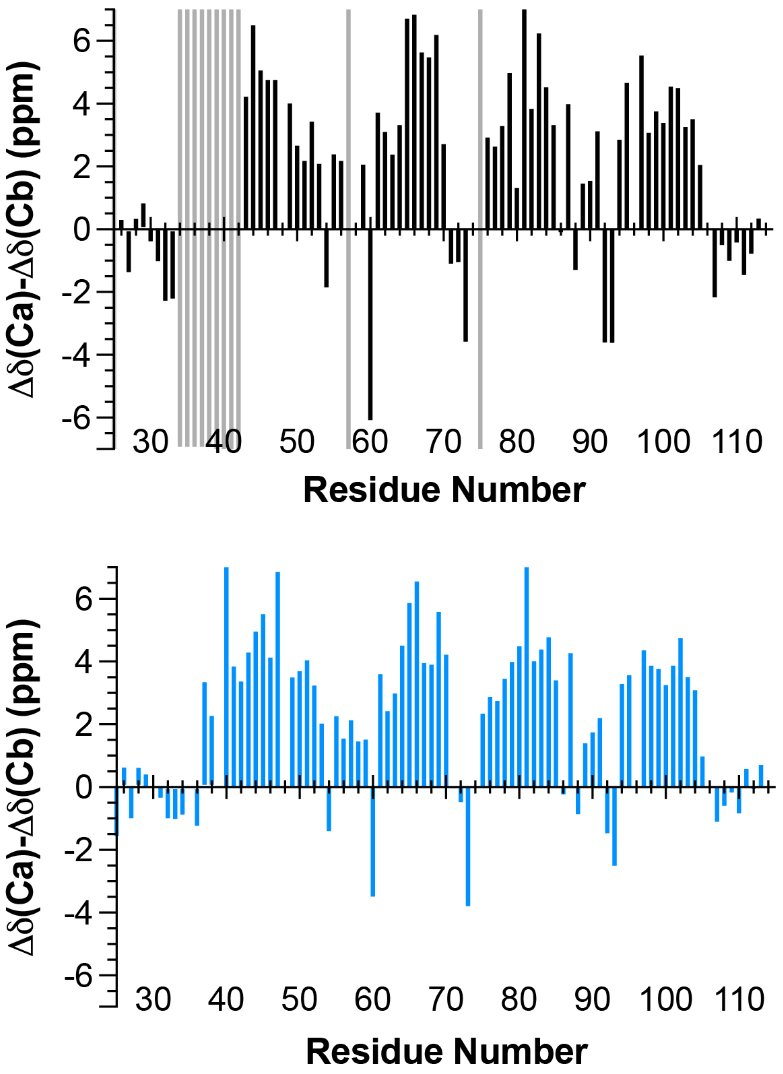


**Figure S2. Experimental and AlphaFold-predicted secondary chemical shifts of RTL8C ΔN show good agreement.** Experimentally determined $C_\alpha$-$C_\beta$ values (top) compared against predicted $C_\alpha$-$C_\beta$ values (bottom) from the Alphafold-derived RTL8C structure. Predicted chemical shifts were from EFG-CS [58] (see Methods).

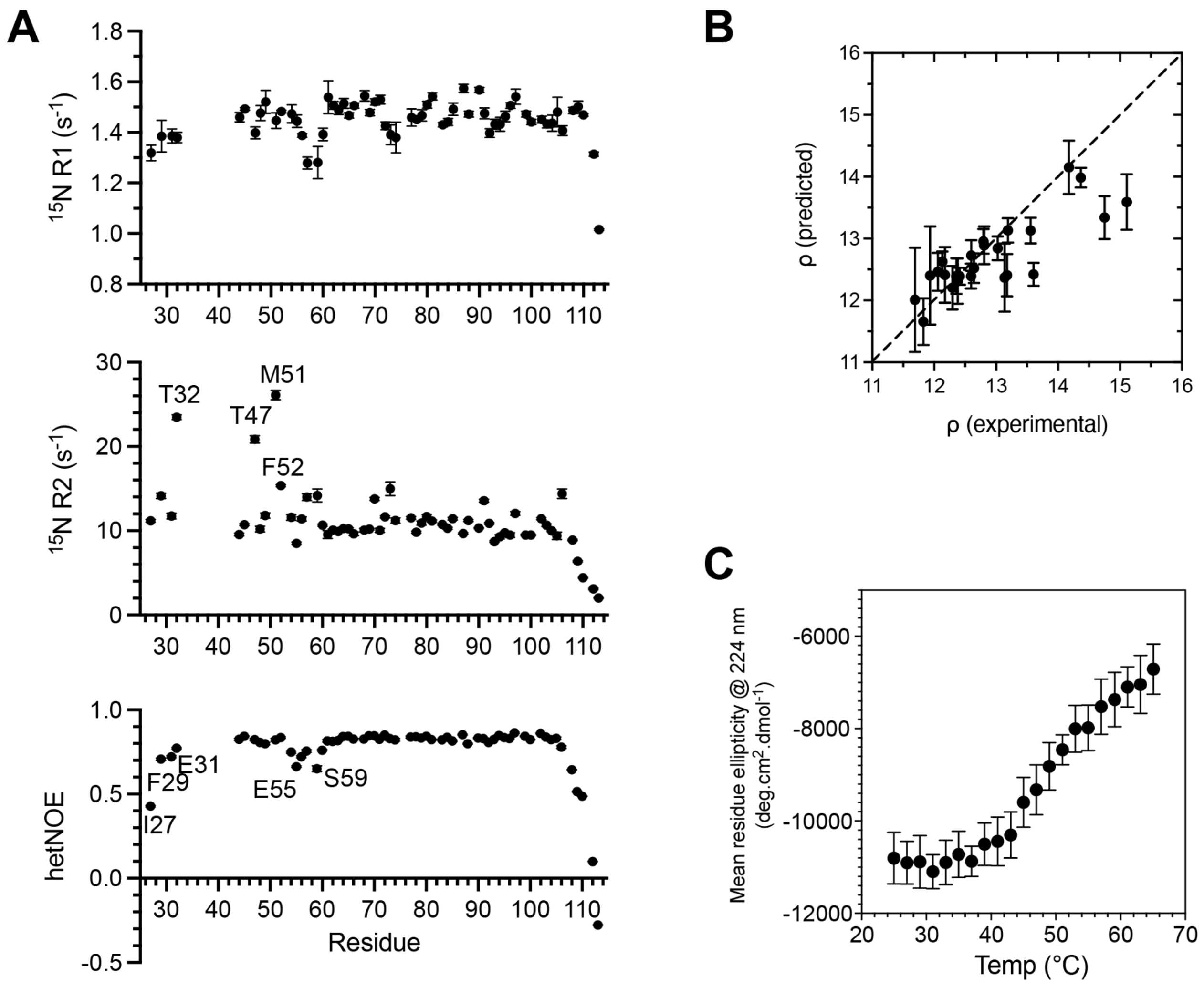


**Figure S3. Biophysical characterization of RTL8C ΔN.** (A) Relaxation data for backbone amide resonances of RTL8C ΔN at pH 6.2 in NaPhosphate buffer at 25°C. Data for $^{15}$N $R_1$ and $R_2$ relaxation rates were collected using 50 µM RTL8C ΔN while {$^1$H-$^{15}$N} steady-state hetNOE values were determined with a 200 µM RTL8C ΔN sample. Errors in $R_1$ and $R_2$ were determined using 500 Monte Carlo trials using RELAXFIT on MATLAB. Errors in hetNOE were determined using the standard error propagation formula. Labeled resonances were colored green in Fig. 2E. (B) The relaxation rates and hetNOE values for rigid elements of secondary structure (see Methods for residue numbers) were used to determine the rotational diffusion tensor of RTL8C ΔN. Plotted are the experimental ρ $(2^*(R_2'/R_1') - 1)^{-1}$ vs. the predicted ρ values from an axially-symmetric diffusion tensor (parameters listed in **Table S1**). From this tensor measurement, rotational correlation time was determined to be 5.9 ns, consistent with the expected molecular weight (10.4 kDa) of RTL8C ΔN [68]. (C) Secondary structure of 5 µM RTL8C ΔN was monitored by far-UV CD signal at 224 nm as a function of temperature. Error bars represent the instrument-reported error averaged over a 15 s acquisition window at each temperature.

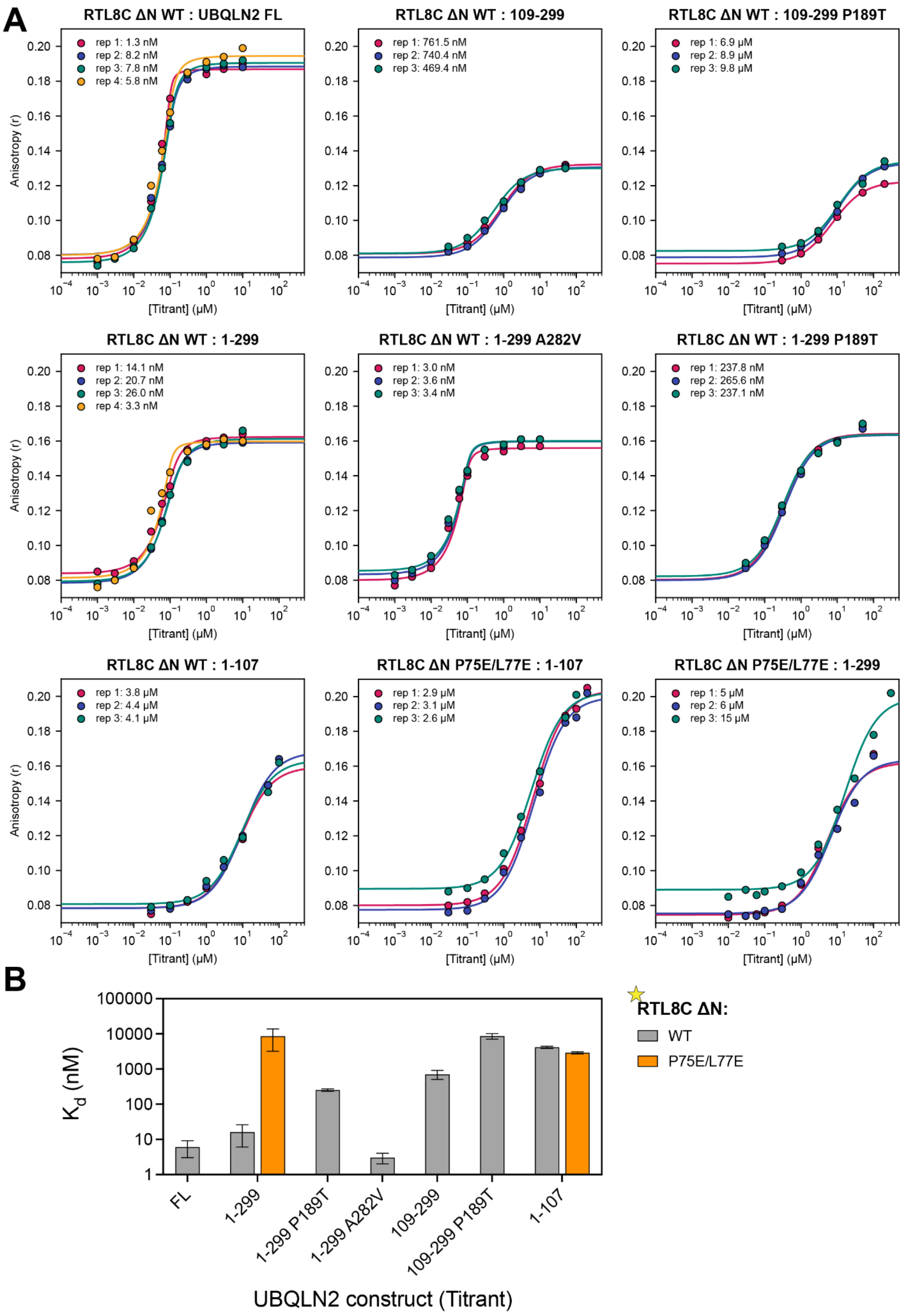


**Figure S4. Summary of RTL8C ΔN:UBQLN2 binding affinities.** (A) Individual FA binding curves for AF488 labeled RTL8C ΔN (WT or P75E/L77E) titrated with UBQLN2 constructs. Each replicate was fit independently using a single-site binding model. (B) Summary of equilibrium dissociation constants ($K_d$) from the fits in panel A. Values are mean ± SD from replicates.

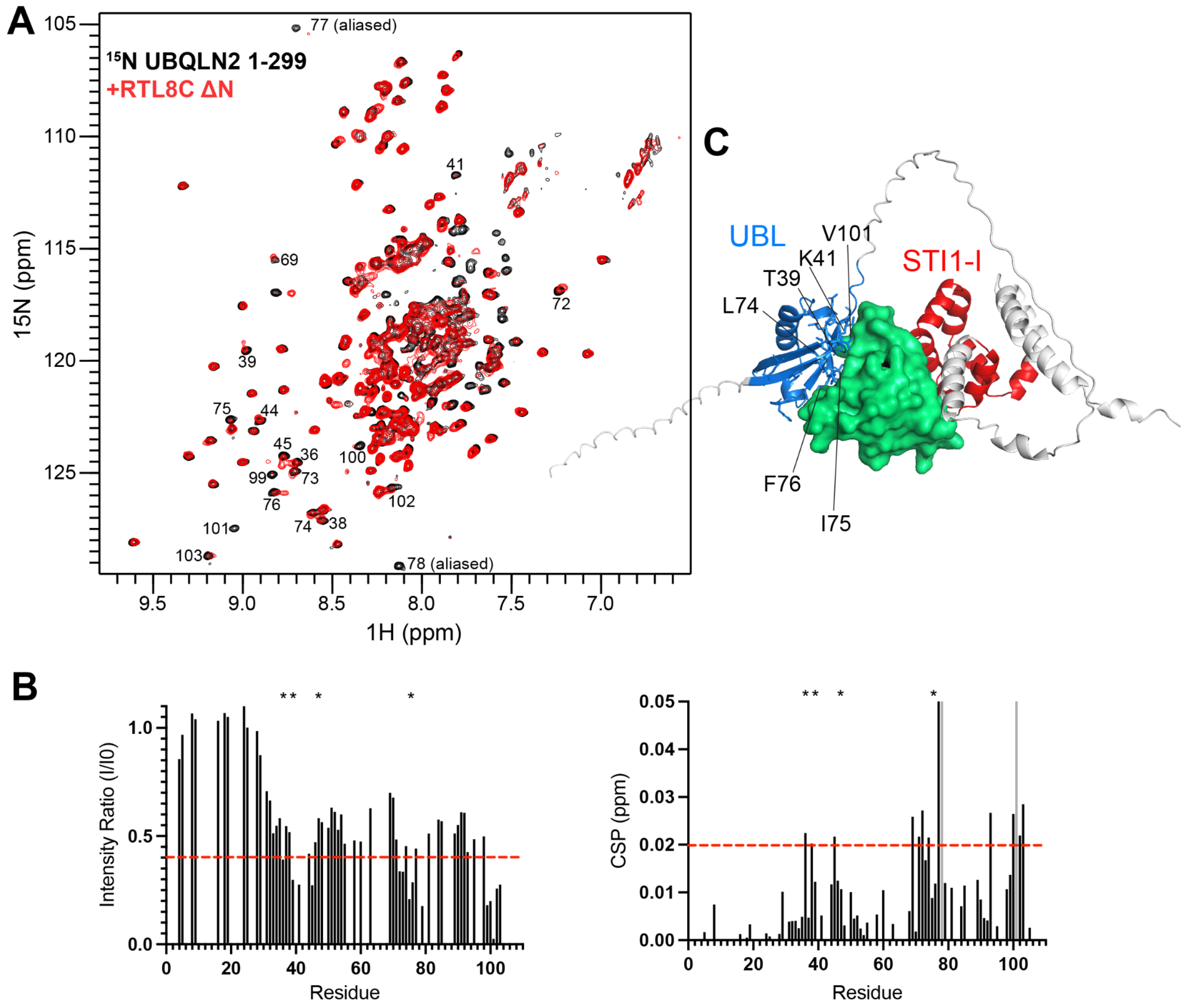


**Figure S5. NMR mapping of the RTL8C ΔN interaction surface on the UBQLN2.** (A) $^{15}N$-$^{1}H$ TROSY-HSQC spectrum of 50 μM UBQLN2 1-299 at 40 ºC (pH 6.2 NaPhosphate buffer) in the absence (black) and presence (red) of equimolar amount of RTL8C ΔN. Amide resonances with either significant signal attenuations or chemical shift perturbations (CSPs) are annotated. (B) Intensity ratio of UBQLN2 UBL (residues 33-107) amide resonances (left) and CSPs (right) with and without RTL8C ΔN. Asterisks mark resonances with two signals (potential slow exchange on the NMR timescale). (C) Intensity ratios below 0.4 and CSPs greater than 0.02 ppm (red dashed lines in panel B) are mapped onto the predicted UBQLN2 1-299:RTL8 ΔN complex and are marked as blue sticks.

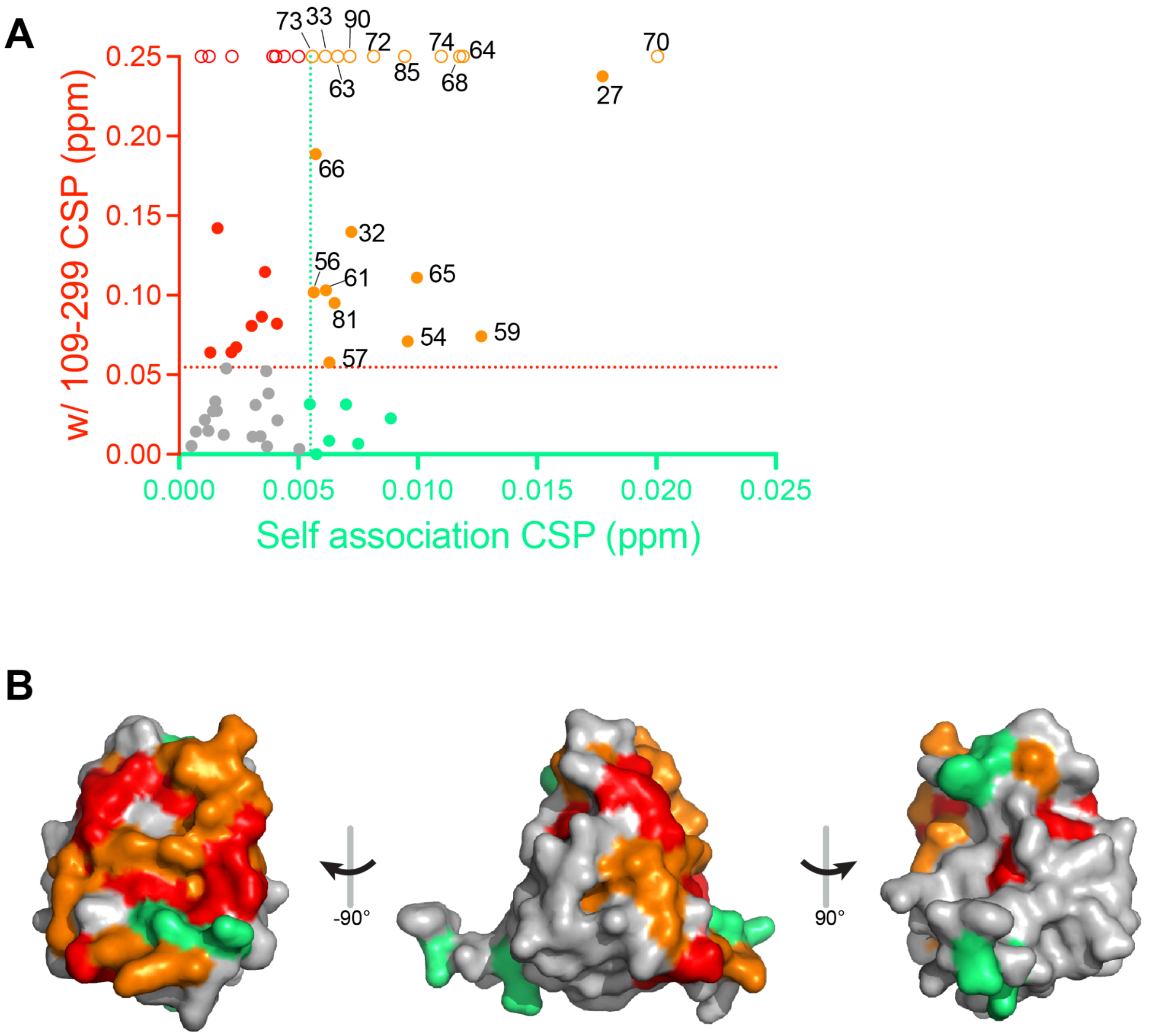


**Figure S6. RTL8C ΔN self-association surface overlaps with the UBQLN2 STI1-I binding surface.** (A) Per-residue CSP correlation from $^{15}$N RTL8C ΔN self-association (50 vs. 200 μM; x-axis, **Fig. 2A**) and 50 μM $^{15}$N RTL8C ΔN titrated with UBQLN2 109-299 (y-axis, **Fig. 4B**). Dashed lines mark the mean CSP of each dataset (thresholds for perturbation). Residues exceeding thresholds are shown: for both datasets in orange, for 109-299 only in red, and for self-association only in green; neither in gray. Open circles: RTL8C ΔN peaks broadened beyond detection in the presence of UBQLN2 109-299 (assigned arbitrary y value of 0.25). (B) Surface representation of the RTL8C ΔN AlphaFold model (AF-A6ZKI3; residues 25-113 are shown), colored by the same scheme as in panel A. Common perturbed residues (orange) cluster on one contiguous surface.

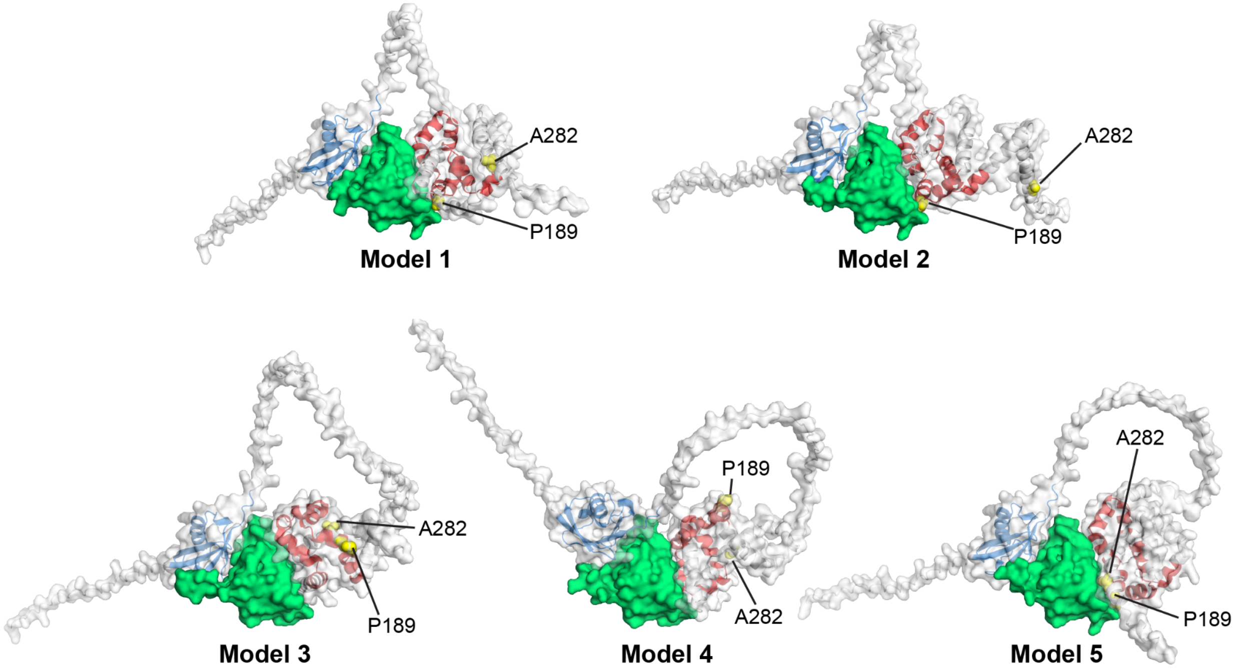


**Figure S7**. **AlphaFold-Multimer models of the UBQLN2 1-299:RTL8C ΔN complex.** AlphaFold predictions of UBQLN2 1-299 in complex with RTL8C ΔN, shown as five independent models (model 1, top, to model 5, bottom). UBQLN2 1-299 in surface/cartoon representation; RTL8C ΔN in green surface representation. Within UBQLN2: Marine blue is UBL domain (residues 33-107), and red is STI1-I domain (residues 178-247). P189 and A282 are labeled (yellow spheres).

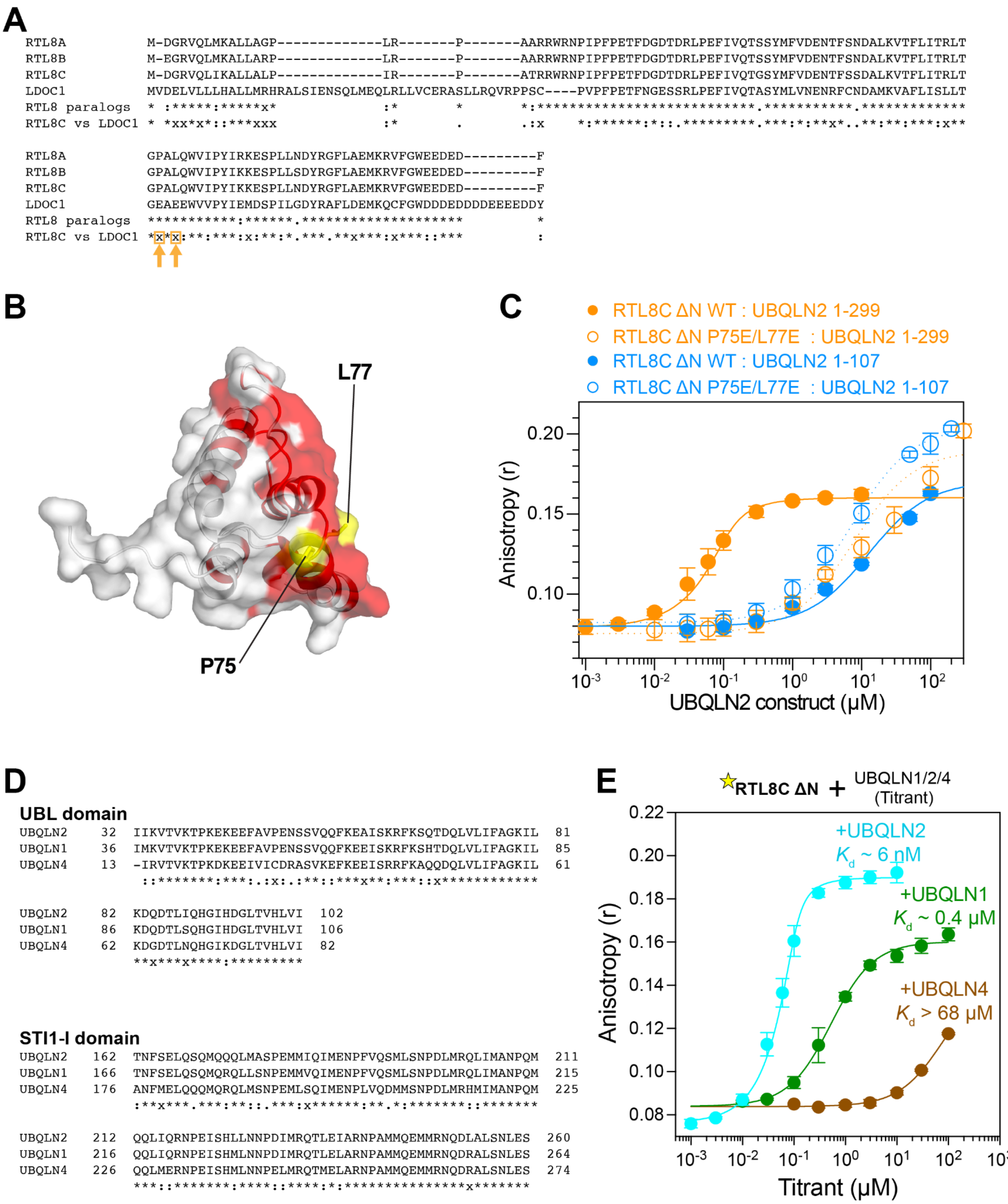


**Figure S8. The selectivity of UBQLN2-RTL8 binding does not extend to their paralogs.** (A) Multiple sequence alignment of RTL8 paralogs and LDOC1. Conservation tracks are shown (between RTL8 paralogs and RTL8C vs LDOC1) as identical (*), strongly conserved (:), weakly conserved (.), and non-conserved (x); gap/indels are blank. Orange arrows in the RTL8C vs LDOC1 track mark P75E/L77E substitutions used in next panel. (B) P75 and L77 positions (yellow) are mapped on AlphaFold-predicted structure of RTL8C ΔN (AF-A6ZKI3-F1-v6). UBQLN2 STI1-I binding surface of RTL8C ΔN is shown in red (as in Fig. 4C). (C) FA UBQLN2 1-299 and 1-107 titration curves compared between WT and P75E/L77E AF488 labeled RTL8C ΔN. Data are mean ± standard deviation (SD); n≥3. The mutations do not affect UBL engagement, as P75E/L77E binds 1-299 or 1-107 with affinities comparable to WT RTL8C ΔN: UBQLN2 1-107 (**Table 1**; **Fig. S4**), indicating that the substitutions act locally at the STI1-I interface rather than globally disrupting the protein. (D) Sequence alignments of the UBL and STI1-I domains of UBQLN1, UBQLN2 and UBQLN4. Conservation below the alignment. (E) Fluorescence anisotropy experiments of titrating UBQLN1, UBQLN2, and UBQLN4 constructs to AF488-labeled RTL8C ΔN. Total RTL8C ΔN concentration was 100 nM.

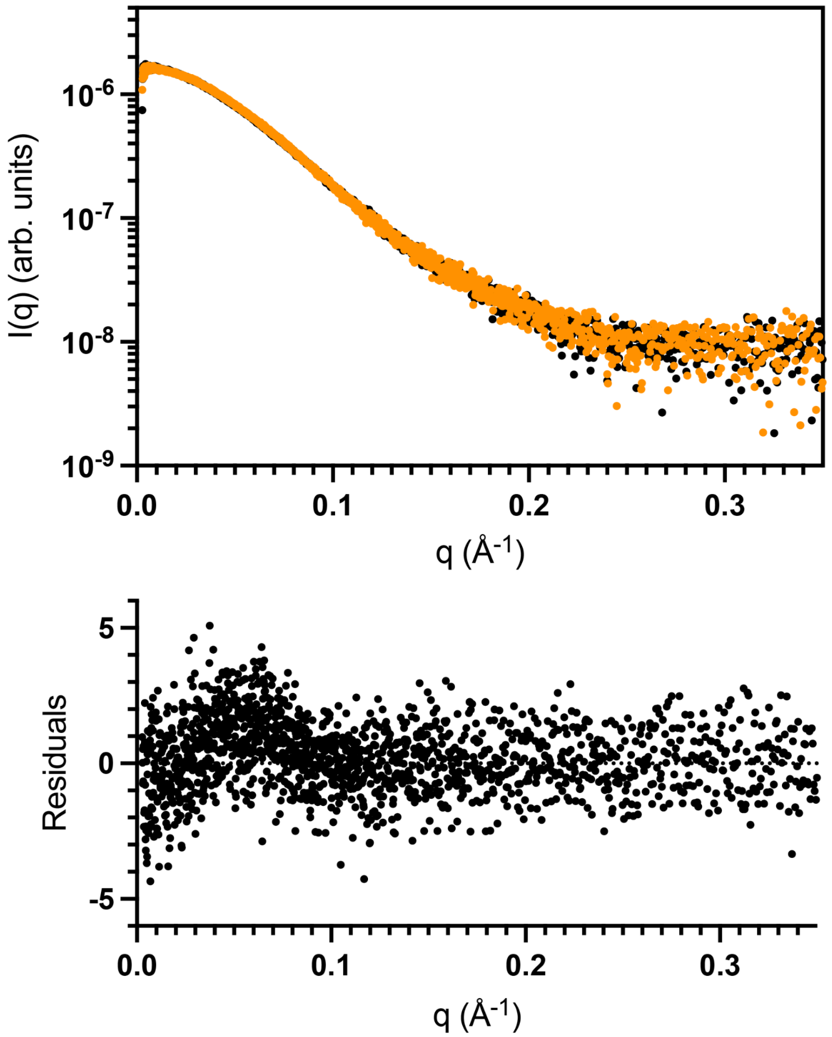


**Figure S9. Comparison of SAXS profiles** for the 1:1 RTL8C ΔN:UBQLN2 1-299 complex at 20 ºC (black) and 40 ºC (orange) with normalized residuals ( shown below the SAXS profile.

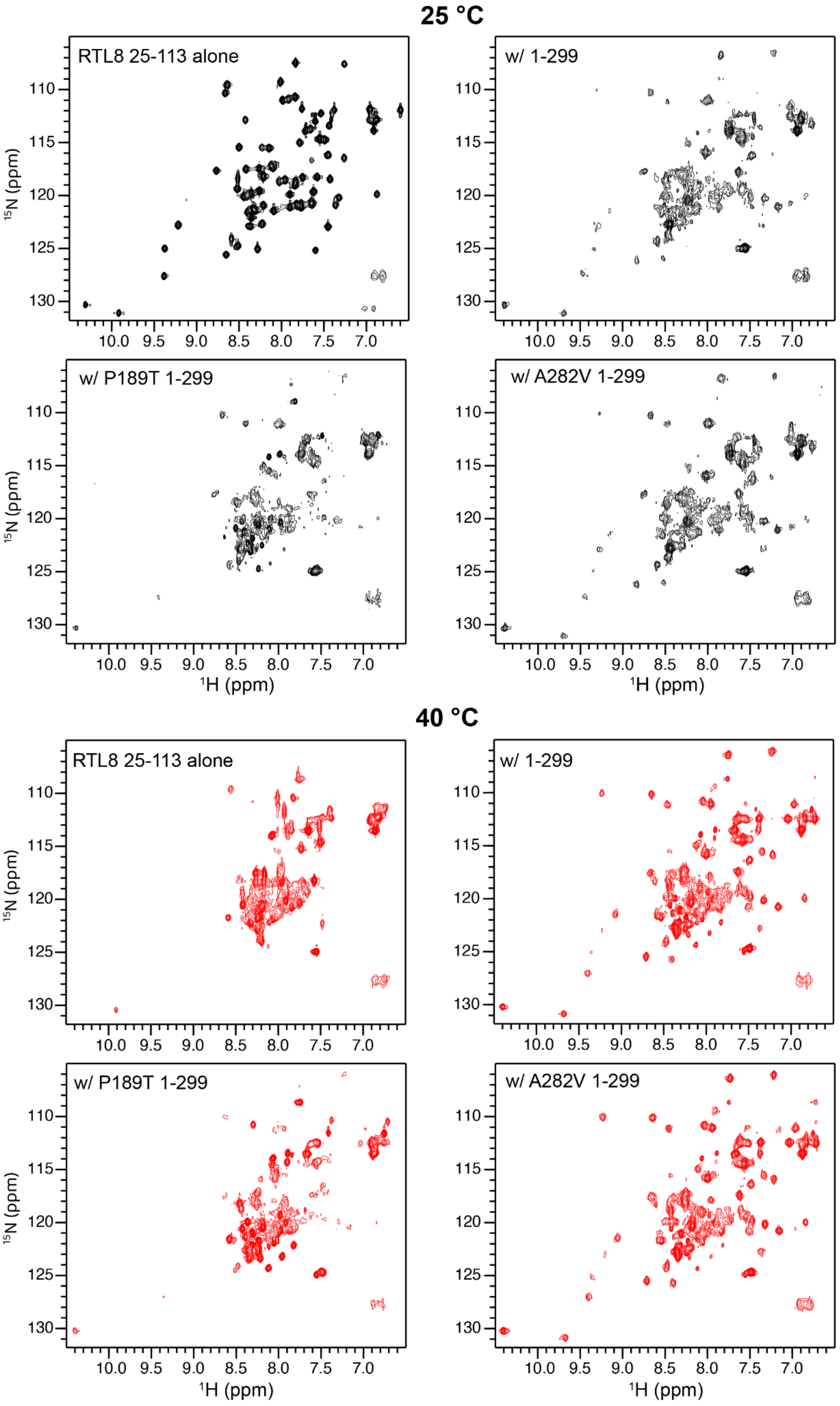

25 °C
RTL8 25-113 alone
w/ 1-299
w/ P189T 1-299
w/ A282V 1-299
40 °C
RTL8 25-113 alone
w/ 1-299
w/ P189T 1-299
w/ A282V 1-299
15N (ppm)
1H (ppm)
110
115
120
125
130
10.0
9.5
9.0
8.5
8.0
7.5
7.0

**Figure S10. NMR spectra of RTL8C ΔN in the presence of WT or mutant UBQLN2 1-299 at 25°C and 40°C.** $^{1}H$-$^{15}N$ SOFAST-HMQC NMR spectra of RTL8C ΔN either alone (50 μM) or in the presence of equimolar amounts of UBQLN2 1-299 variants, specifically WT, P189T, or A282V all in 20 mM HEPES buffer at pH 7. At 25 ºC, all acquisition and processing parameters are the same with the exception that RTL8 25-113 has 2x higher base contour level compared to the other spectra. At 40°C, all acquisition and processing parameters are the same but the base contour level is slightly different than the set of spectra at 25°C.

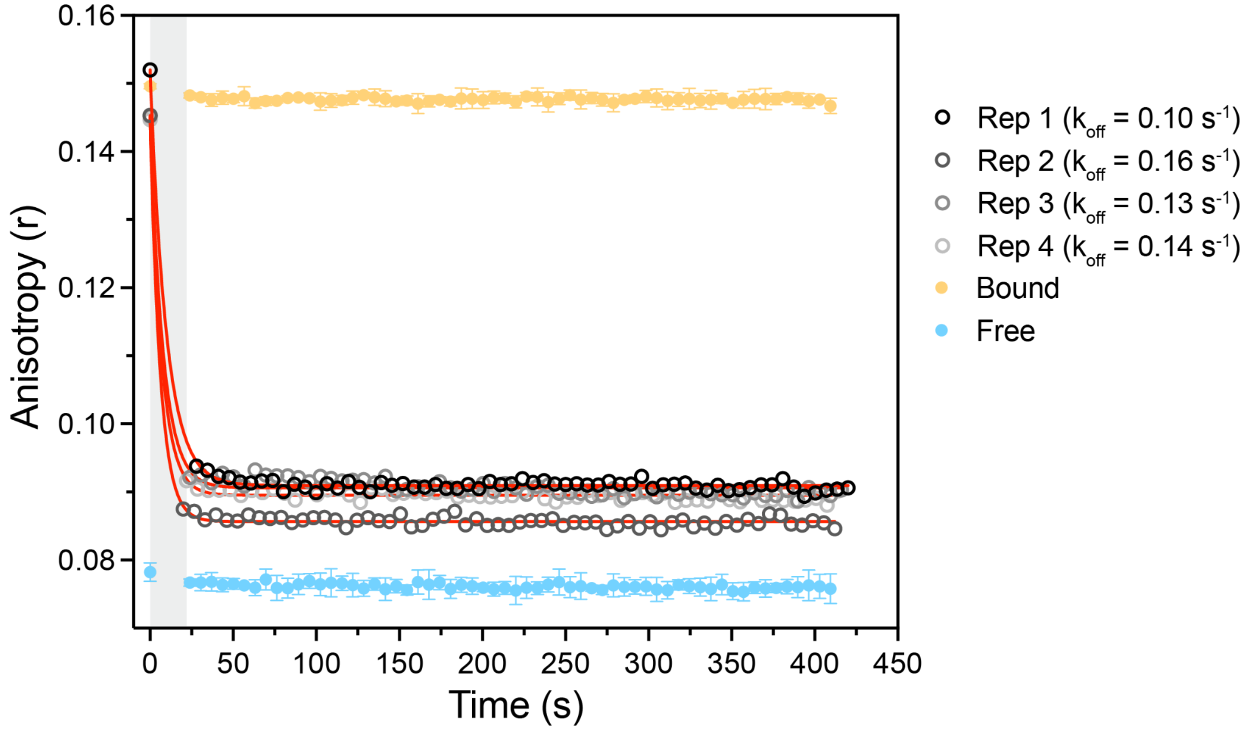


**Figure S11. Dissociation kinetics of the RTL8C ΔN:UBQLN2 1-299 complex measured by fluorescence anisotropy chase assay.** The pre-formed complex of AF488-labeled RTL8C ΔN (50 nM) and unlabeled UBQLN2 1-299 (300 nM) was chased with unlabeled RTL8C ΔN (15 μM, 50-fold molar excess over UBQLN2) at 25 °C. Anisotropy decay, corrected for the dead time (gray shaded region 20-28 s) and instrumental timing offset (see Methods), was fit to a single-exponential decay model (values in black and fit in red) yielding $k_{off}$ ~ 0.13 $s^{-1}$; t½ ~ 5.3 s. As dead time was comparable to the fitted t½, reported $k_{off}$ values represent lower-limit estimates of the true dissociation rate. Free AF488-RTL8C ΔN (50 nM) and the bound complex (with 300 nM UBQLN2 1–299) served as separate controls. Notably, anisotropy after chase addition does not fully return to the free RTL8C ΔN baseline for 50 nM, consistent with RTL8C ΔN self-association at the chase concentration used (15 μM; see **Fig. 2F**).

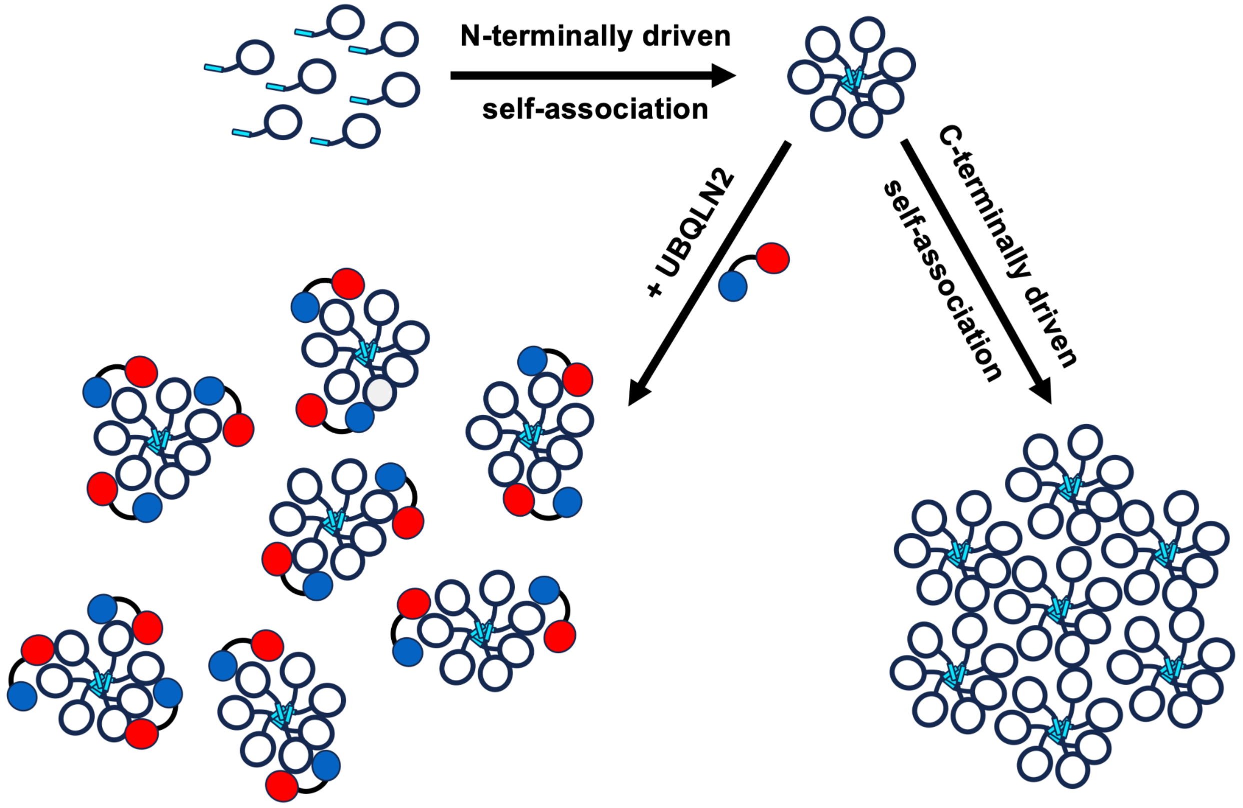


**Figure S12. A model of RTL8 self-association.** RTL8 exists as high-order oligomers via self-association of its N-terminus (residues 1-24). In the absence of UBQLN2, the RTL8 C-terminus (residues 25-113 or RTL8 ΔN) self-associates, leading to even larger oligomers (possibly aggregates) that need to be removed. In the presence of UBQLN2 (construct shown is UBQLN2 1-299 with blue and red representing the UBL and STI1-I domains, respectively), the N-terminally-driven RTL8 oligomers stay separated with the C-terminus exposed to bind to UBQLN2 and other binders such as PEG10.

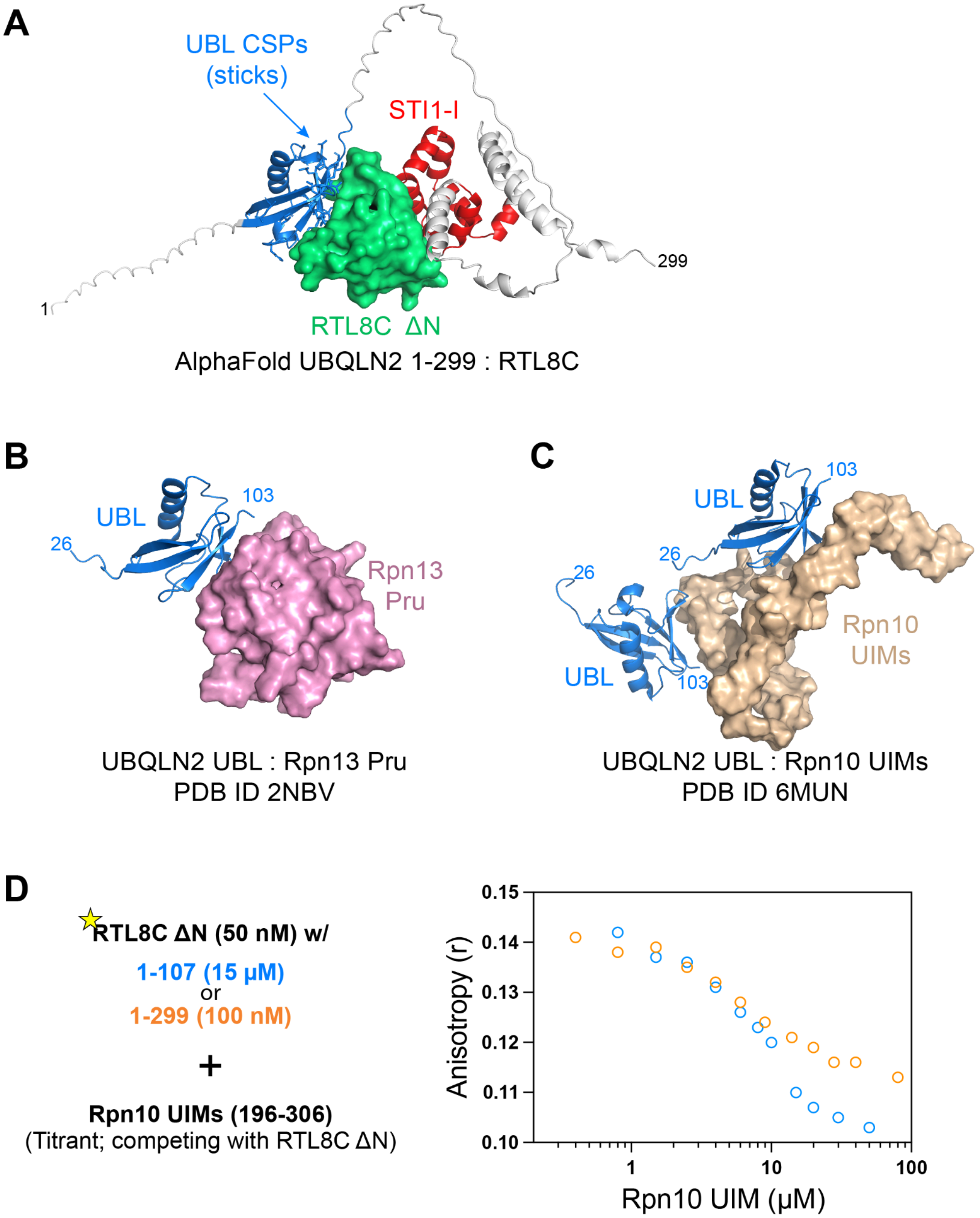


**Figure S13. Comparison of AlphaFold predicted UBQLN2:RTL8C ΔN complex with UBL:Rpn solution-derived structures.** (A-C) Note that the UBL binding surface for the complex in panel A (blue sticks, same as in **Fig. S5** above) matches the same UBL binding surface when interacting with Rpn13 Pru domain (panel B) or Rpn10 UIMs (panel C). The UBL domains in all structures were placed in similar orientation for this figure. UBL residues (33-103) and STI1-I residues (178-247) were color-coded blue and red, respectively. RTL8C ΔN (green), Rpn13 Pru (pink), Rpn10 UIMs (wheat) are shown in surface representations. (D) Fluorescence anisotropy-based competition assay between RTL8C ΔN and Rpn10 UIMs (residues 196-306) for the UBQLN2 UBL surface. Schematic: AF488-RTL8C ΔN (50 nM) is pre-bound to a UBQLN2 construct (50-80% bound); titrating in Rpn10 196-306 competes off labeled RTL8C ΔN, decreasing anisotropy.

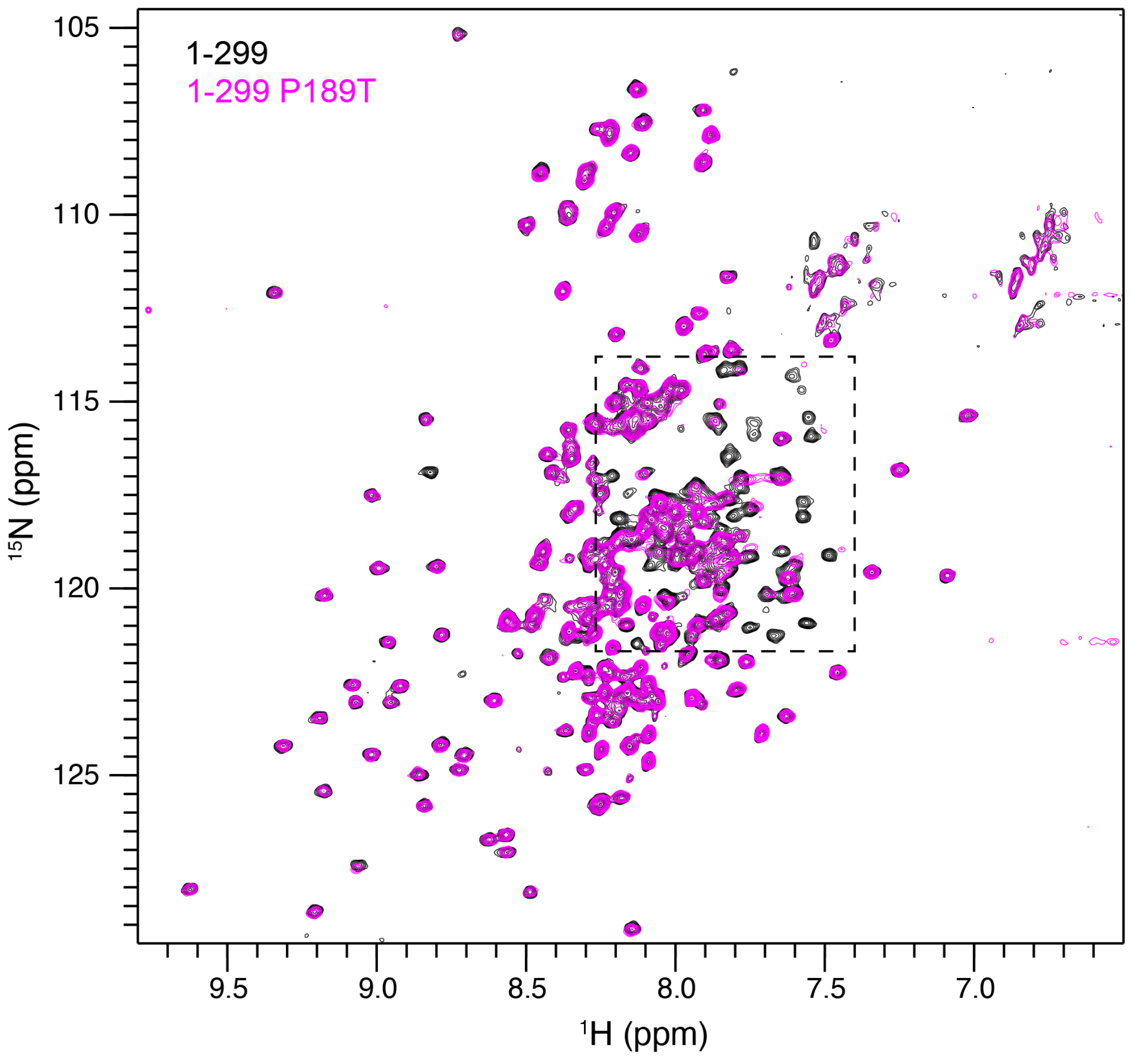


**Figure S14. Comparison of wild-type and mutant UBQLN2 NMR spectra.** $^{15}$N-$^{1}$H TROSY-HSQC spectra of 50 µM UBQLN2 1-299 (black) and 50 µM of UBQLN2 1-299 P189T (magenta) at 40 ºC (pH 6.2 NaPhosphate buffer). Amide resonances generally overlap across the entire spectrum except for the dashed region that corresponds to STI1-I resonances.

**Table S1. Rotational Diffusion Tensor Determination**

| | D⊥ (perpendicular) [a] | D // (parallel) | α [b] | β | $\tau_c$ (ns) [c] | Probability [d] |
|---|---|---|---|---|---|---|
| RTL8C ΔN | 2.62 (0.13) | 3.30 (0.33) | 161 (29) | 85 (8) | 5.86 (0.07) | $1.73 \times 10^{-8}$ (ax/iso) |

[a] Principal values of the diffusion tensor, in $10^7$ $s^{-1}$. Errors (in parentheses) were estimated using 1000 Monte Carlo trials.
[b] Euler angles (in degrees) characterize the principal axes frame of the diffusion tensor with respect to the PDB coordinate frame of the AlphaFold-predicted structure of RTL8C ΔN.
[c] Overall rotation correlation time.
[d] F-test-derived probability that the improvement in the fit for an axially symmetric diffusion tensor over the isotropic diffusion tensor models could have occurred by chance.

**Table S2. SEC-MALS-SAXS data at 20 °C and 40 °C**

| | Temperature (ºC) | Expected MW (kDa) | Observed MW (MALS) (kDa) ($M_{avg}$) | Guinier $R_g$ (Å) [a] | GNOM $R_g$ (Å) [b] |
|---|---|---|---|---|---|
| RTL8C FL | 20 | 13 | $1.158 \times 10^3$ | | |
| RTL8C ΔN | 20 | 10.4 | 10.44 | 14.9 ± 0.2 | 15.0 ± 0.2 |
| RTL8C ΔN | 40 | 10.4 | 20.5 | | |
| UBQLN2 1-299 | 20 | 32.4 | 33.0 | 34.9 ± 0.2 | 36.1 ± 0.3 |
| 1:1 UBQLN2 1-299:RTL8C ΔN | 20 | 42.7 | 40.93 | 29.9 ± 0.1 | 30.9 ± 0.2 |
| 1:1 UBQLN2 1-299:RTL8C ΔN | 40 | 42.7 | 40.52 | | |

[a] $R_g$ and standard errors were determined from the linear fit of ln(I(q)) vs $q^2$.
[b] $R_g$ and standard errors determined using GNOM [52].

**Table S3. Results of CALVADOS simulations of UBQLN2 1-299 bound to RTL8C ΔN**

| Construct | Simulated $R_g$ (Å) | Domain 1 [a] | Domain 2 [a] | Domain 3 [a] | Additional Constraints |
|---|---|---|---|---|---|
| UBL:RTL8C ΔN:STI1-I | 29.2 ± 0.2 | [32,102] | [162,260] | RTL8C ΔN [25,106] | [[32,102],[162,260], RTL8C ΔN [25,106]] |
| UBL:RTL8C ΔN:UBL bound, STI1-I | 40.3 ± 0.6 | [32,102] | [162,260] | RTL8C ΔN [25,106] | [[32,102], RTL8C ΔN [25,106]] |
| RTL8C ΔN:STI1-I bound, UBL | 36.9 ± 0.2 | [32,102] | [162,260] | RTL8C ΔN [25,106] | [[162,260], RTL8C ΔN [25,106]] |

[a] Folded domain boundaries of the multidomain proteins used in this study. Brackets indicate the first and last residue of the UBQLN2 domain, unless otherwise noted. To keep certain domains together, additional constraints were applied between the listed folded domains (last column). Note that RTL8C ΔN is a separate chain.

**Table S4. SAXS Data Collection Details.**

| | | | |
|---|---|---|---|
| ***(a)* Sample details** | | | |
| Organism | *H. sapiens* | | |
| Source (Catalogue No. or reference) | Expressed in *E. coli* (this work) | | |
| Description: sequence (including Uniprot ID + uncleaved tags), bound ligands/modifications, etc. | RTL8C ΔN (no tags) | UBQLN2 1-299 (no tags), Uniprot ID Q9UHD9 | 1:1 mix of UBQLN2 1-299 (no tags) and RTL8C ΔN (no tags) |
| Extinction coefficient ε in $M^{-1}$ $cm^{-1}$ (wavelength in nm) | 15470 (280) | 2980 (280) | 18450 (280) |
| Molecular mass M from chemical composition (Da) | 10,358 | 32,389 | 42,747 |
| For SEC-SAS, concentration (mg $ml^{-1}$), injection volume (μl), flow rate (ml $min^{-1}$) | 4, 200, 0.6 | 7.41, 200, 0.6 | 5.6, 200, 0.6 |
| Solvent composition and source | pH 6.8 20 mM sodium phosphate | | |
| ***(b)* SAS data collection parameters** | | | |
| Instrument | BioCAT (Sector 18, APS), with Pilatus3 X 1M detector (Dectris) | | |

| | | | |
|---|---|---|---|
| Wavelength (Å) | 1.033 | 1.033 | 1.033 |
| Camera length (m) | 3.702 | 3.702 | 3.702 |
| *q*-measurement range | 0.0032-0.33 | 0.0032-0.33 | 0.0032-0.33 |
| Normalization | Transmitted intensity | Transmitted intensity | Transmitted intensity |
| Exposure time/number | 0.5 seconds (2470 frames) | 0.5 seconds (2470 frames) | 0.5 seconds (2470 frames) |
| Sample Configuration | SEC-MALS-SAXS using a Superdex 200 Increase 10/300 GL column and an Agilent 1260 Series HPLC. UV data was measured with an Agilent 1290 DAD, and MALS/RI data by DAWN HELEOS-II (18-angle) and Optilab T-rEX (RI) instruments (Wyatt Technology). SAXS data was measured in a 1 mm, Mica-windowed flow cell | | |
| Sample Temperature | 20 ºC | 20 ºC | 20 ºC |
| ***(c)* Software employed** | | | |
| SAXS data reduction | Radial averaging; frame comparison, averaging, and subtraction done using BioXTAS RAW 2.4.0 or 2.4.1 | | |
| Basic analysis: Guinier, M.W. P(r) | Guinier fit and M.W. using BioXTAS RAW. RAW uses MoW and Vc M.W. methods | | |
| MALS-RI analysis | Astra 8 (Wyatt) | | |
| ***(d)* Structural Parameters** | | | |
| Guinier Analysis | RTL8C ΔN | UBQLN2 1-299 | 1:1 UBQLN2 1-299:RTL8C ΔN |
| I(0) | 0.0021 | 0.0156 | 0.0185 |
| $R_g$ (Å) | 14.9 ± 0.2 | 34.9 ± 0.2 | 29.9 ± 0.1 |
| q-range (Å$^{-1}$) | 0.00579-0.08622 | 0.00325- 0.02893 | 0.00325-0.03401 |

| | | | |
|---|---|---|---|
| Quality-of-fit parameter (with definition) | 0.8387 ($r^2$) | 0.9460 ($r^2$) | 0.9364 ($r^2$) |
| *M* from MALS (kDa) | 10.44 | 33.0 | 40.93 |

**Supplementary Movies:**

**Movie S1. Ensemble of UBQLN2 1-299 bound to RTL8C ΔN**. Reconstructed ensemble of UBQLN2 1-299 bound to RTL8C ΔN using structural constraints delineated in **Table S3** for UBL:RTL8C ΔN:STI1-I. Colors are blue (UBL), red (STI1-I), dark grey (IDRs in UBQLN2), and white (RTL8C). The movie frame rate corresponds to 100 ps per second.
